# Non-convergent aridity adaptation despite pervasive linked selection in *Eucalyptus*

**DOI:** 10.64898/2026.08.25.747154

**Authors:** Maddie E. James, Travis G. Britton, Jonathan D. Mitchell, Ben Halliwell, Barbara Holland, Ian J. Wright, Daniel Ortiz-Barrientos

## Abstract

Whether independent lineages evolve similar genetic solutions when faced with the same environmental pressure is central to understanding how repeatable and predictable adaptation is. Answering this question is increasingly urgent as climate change intensifies drought and aridity worldwide, a shared pressure to which many species must independently adapt. Here we characterised genomic adaptation in *Eucalyptus* across three independent species pairs, each comprising two closely related lineages that have diverged from wetter into drier environments. Across all pairs we found concordant genome-wide landscapes of diversity and divergence, and strong evidence for pervasive linked selection. Because linked selection acts most strongly in the same conserved features of the genome, this shared architecture could concentrate differentiation in the same regions across lineages, creating an appearance of convergent adaptation. Despite this, the genomic regions associated with the transition to drier environments were largely non-convergent, with almost no sharing of outlier loci among pairs, demonstrating that each lineage adapted through a largely independent genetic route. Some convergence was instead evident at the level of biological function, indicating that lineages reached similar functional outcomes using different genes. We further identified large genomic islands of differentiation, which were dominated by the signature of linked selection rather than elevated divergence, though several harboured candidate genes within the drought and abscisic acid regulatory networks. Together, our results indicate that adaptation to aridity in *Eucalyptus* is complex and polygenic, and largely unpredictable at the level of individual loci.

## INTRODUCTION

A fundamental question in evolutionary biology is whether adaptation is predictable. For instance, when independent lineages face the same environmental challenges, do they follow the same adaptive trajectories? Is the underlying genetic architecture of adaptation relatively simple (involving few genes) or more complex (involving many)? Answering these questions in natural systems is challenging, as it is difficult to associate adaptive traits with their underlying genes (Barrett & Hoekstra, 2011; Hoban et al., 2016; Mackay et al., 2009; Ravinet et al., 2017; Rockman, 2012). One powerful approach is to study systems of convergent evolution (also known as parallel, replicated or repeated evolution) (Cerca, 2023; James et al., 2023; Schluter & Nagel, 1995). Throughout this paper, we use the term *convergent* to describe cases where distantly related lineages evolve similar phenotypes in response to similar environmental conditions, and *parallel* for closely related lineages. Such systems can be viewed as ‘natural replicates’ of an evolutionary solution (Lenormand et al., 2009). When the same genomic regions harbour signatures of selection across independent replicates, these repeated signatures are unlikely to have arisen by chance. This suggests that natural selection, rather than genetic drift, underlies their evolution (Losos, 2011; Orr, 2005; Schluter & Nagel, 1995). Examining such systems allows us to ask not only which genes underlie adaptation, but how repeatable and predictable the genetic basis of adaptation is.

The most well-documented examples of genetic convergence and parallelism involve simple genetic architectures, where the same or a few genes are recruited repeatedly across independent lineages. For example, in the classic three-spine stickleback system, replicate freshwater populations have independently evolved reduced pelvic armour through repeated selection on alleles at the Eda gene (Colosimo et al., 2005). Such cases of simple genetic architectures suggest that evolution can be highly predictable at the molecular level (Conte et al., 2012; Rosenblum et al., 2014; Stern, 2013; Stern & Orgogozo, 2009; Yeaman et al., 2018). However, for complex quantitative traits controlled by many genes, increasing evidence suggests that adaptation often follows largely unique genetic trajectories even when phenotypic outcomes converge, with different loci contributing to similar traits across independent lineages (Hoitinga & Birkeland, 2025). Interestingly, even when replicate lineages recruit unique sets of genes, the underlying biological functions are often shared (e.g., James et al., 2021; Szukala et al., 2023; Tenaillon et al., 2012; Yeaman et al., 2016), highlighting that the repeatability of the genetic basis of adaptation can manifest at different levels of biological organisation (Allard & Kumar, 2026). Yet, how repeatable the genetic basis of adaptation is, and at what level of biological organisation, remains unclear.

The genomic landscape of divergence is not only shaped by natural selection acting on adaptive loci, but also by the way selection interacts with recombination across the genome. For instance, if a selective sweep arises in a region of low recombination, a swath of linked sites will hitchhike with the selected allele and reduce diversity in the region, and this reduction will extend over a greater physical distance than in regions of high recombination (Begun & Aquadro, 1992; Cutter & Payseur, 2013). Selection against deleterious alleles, or background selection, produces a similar positive correlation between diversity and recombination (Charlesworth et al., 1993). Together, these processes are known as linked selection, and their strength varies across the genome according to the local recombination rate and the density of sites under selection.

Linked selection appears to be a pervasive feature of many genomes, and has been shown to shape broad patterns of diversity and differentiation in a wide range of taxa including *Drosophila*, birds, and plants (Burri et al., 2015; Elyashiv et al., 2016; Liang et al., 2022). Importantly, because linked selection acts most strongly in regions of low recombination and high gene density, independent lineages that share a similar genome architecture can evolve similar differentiation landscapes, with the same genomic regions repeatedly emerging as differentiation peaks rather than reflecting repeated adaptation (Burri, 2017; Chase et al., 2021; Cruickshank & Hahn, 2014). This has important implications for studies of convergent and parallel evolution, as apparent repeatability in the genomic basis of adaptation may partly reflect the shared action of linked selection rather than the reuse of the same adaptive loci. Characterising the genome-wide relationship between differentiation and diversity is therefore an important step to help identify candidate regions of repeated adaptation.

Eucalypts are one of Australia’s most widespread and iconic plant taxa, occurring in every major biome in Australia and displaying a wide range of phenotypes (Bennett, 2016). Radiation of *Eucalyptus* into arid environments has occurred relatively recently, with rapid drying of the Australian continent since the Pliocene (5.3 Mya) (Thornhill et al., 2019). This group is therefore an ideal system to examine the repeatability of the genetic basis of climatic adaptation, as many clades have independently radiated into harsh environments in the recent past. The repeated adaptation of *Eucalyptus* into arid environments has been associated with shifts in multiple traits (Halliwell et al., 2025), including thicker, high leaf mass per area leaves (Cernusak et al., 2011; de Boer et al., 2016), higher leaf nitrogen per unit area (Turner et al., 2008), greater allocation to stem cross-sectional area per unit leaf area (Anderegg et al., 2021), and denser wood (Pfautsch et al., 2016). Many of these trends have also been detected in common gardens both within-and among-species (Britton et al., 2026; Pritzkow et al., 2020; Schulze et al., 2006; Warren et al., 2006), suggesting that their relationships to aridity likely reflect a genetic basis.

Previous genomic work has begun to reveal the genetic basis of climate adaptation in *Eucalyptus*. Genome-wide scans have shown that signatures of aridity adaptation are distributed broadly across the genome, highlighting a highly complex and polygenic architecture where adaptation involves many loci of small effect (Butler et al., 2022; Filipe et al., 2022; Jordan et al., 2017; Steane et al., 2014, 2017; von Takach et al., 2021). While some climate-associated adaptive signals are shared among species, a substantial component of climate adaptation appears to be lineage specific, with limited sharing of adaptive loci across lineages (Ahrens et al., 2025; Steane et al., 2017). Adding further complexity, different components of drought tolerance are genetically modular. For instance, drought resistance, recovery, and growth are underpinned by largely non-overlapping genomic regions, indicating that distinct aspects of drought response can evolve semi independently (Ammitzboll et al., 2020). Despite this growing body of work, it remains unclear how divergent adaptive trajectories arise across lineages, and how genome wide processes such as linked selection and reduced recombination shape patterns of genomic differentiation during repeated transitions into arid environments.

Here, we characterise the genomic landscape of divergence associated with the transition to aridity across multiple independent *Eucalyptus* species pairs. Each pair consists of two closely related species that differ substantially in the mean annual precipitation across their geographic range and that are connected by an evolutionary transition from wet to drier environments. Specifically, we: (1) investigate the role of linked selection in shaping the genomic landscape of divergence, (2) identify genomic regions showing signatures of divergence associated with the transition to aridity and test whether they are convergent across independent species pairs, (3) characterise the biological pathways underlying these regions, (4) examine how they relate to genomic features such as recombination rate, gene density and GC content, and (5) characterise broad islands of divergence across the genome. Our work sheds further light on whether the genomic basis of adaptation to aridity in *Eucalyptus* is predictable and repeatable or whether independent lineages have followed largely unique evolutionary trajectories.

## METHODS

### Species selection

A published *Eucalyptus* phylogeny (Thornhill et al., 2019) was used to construct four phylogenetically independent precipitation contrasts (also referred to as species pairs, Figure 1B). Each pair consists of two closely related species that differ substantially in the mean annual precipitation (MAP) of their current-day geographical distributions, but experience similar mean annual temperatures (MAT) (Figure 1A,C; Table S1). This design allowed us to reduce the effect of temperature sufficiently that it could be ignored. By examining multiple transitions to aridity, we can determine whether evolution has repeatedly favoured similar genetic solutions to water limitation, or whether each drier lineage has followed a unique adaptive trajectory.

**Figure 1.**
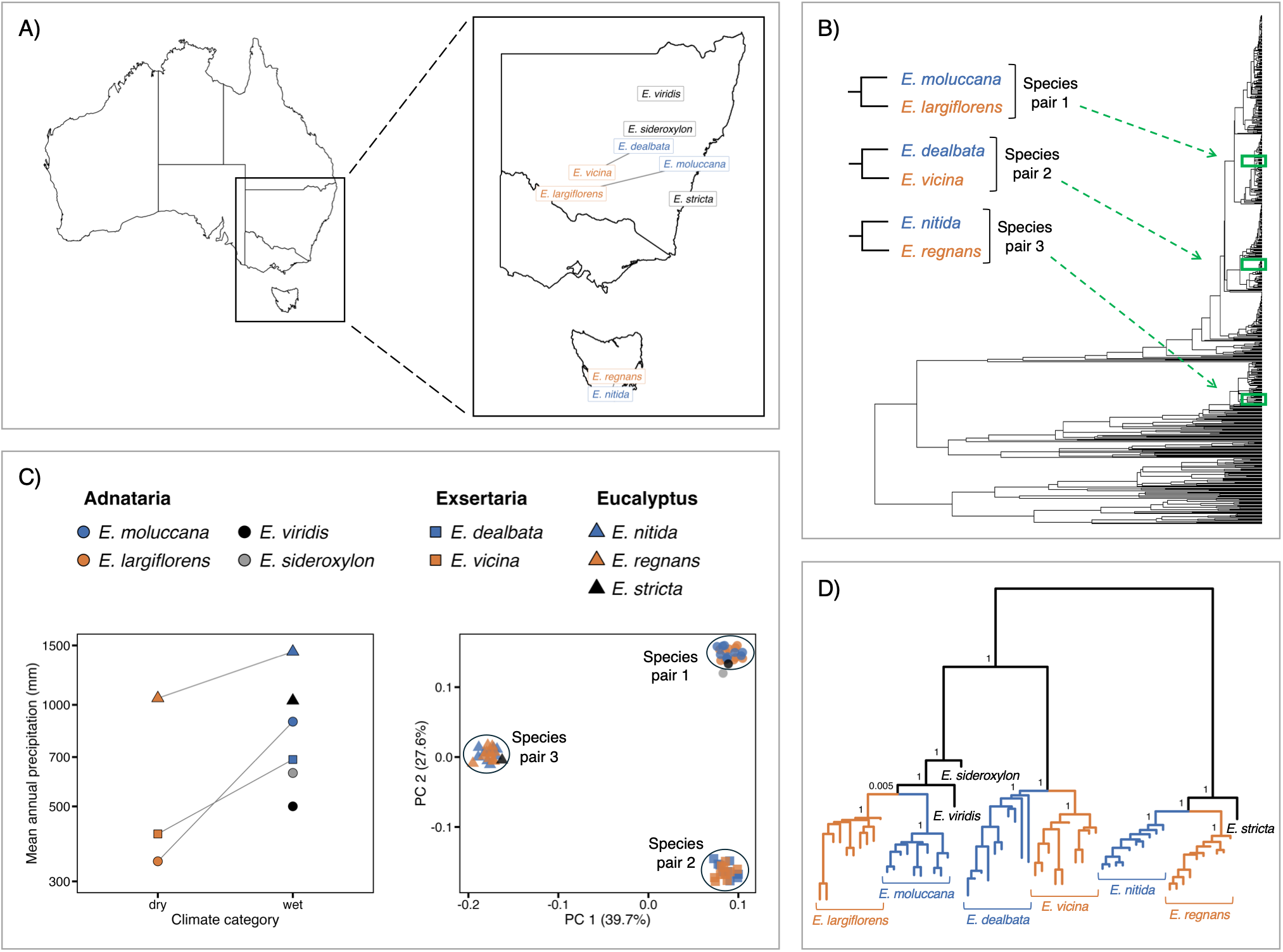
*Eucalyptus* study system and sampling design. A) Map of Australia showing the sampling locations of each species in Australia. B) Time-calibrated *Eucalyptus* phylogeny, modified from Thornhill et al. (2019), denoting the three replicate species pairs of this study, each comprising a wet-adapted lineage (blue) and a drier-adapted lineage (orange). C) Left: mean annual precipitation (mm) of each species across climate categories (dry vs. wet). Species pairs are connected by a line. Points are shaped by taxonomic section – Adnataria (circle), Exsertaria (square) and Eucalyptus (triangle). Right: principal component analysis of 428,330 unlinked SNPs. D) Coalescent-based species tree inferred with ASTRAL-IV. Node values are local posterior probabilities.

Climate variables for all species were determined using previously published ‘climate envelopes’ (Gallagher et al., 2019) based on species’ geographic distributions determined from occurrence records in the Australasian Virtual Herbarium (http://avh.chah.org.au/). MAP (mm) and MAT (°C) were extracted for each grid cell that a species occurred in at a 1 km resolution, using CHELSA climate data (Karger et al., 2017). Mean values for each climate variable were then calculated across all occupied cells for each species. Following standard convention (Singh et al., 2025), we classified species with MAP between 250-500 mm as semi-arid or ‘dry’, and those exceeding 500 mm as ‘wet’. Three species pairs (*E. moluccana–E. largiflorens*, *E. dealbata–E. vicina*, and *E. sideroxylon*–*E. viridis*) represent ‘wet-dry’ contrasts, while one pair (*E. nitida–E. regnans*) represents a ‘wetter-wet’ contrast, enabling us to examine whether adaptive responses depend on absolute aridity thresholds or relative precipitation differences. We also included one additional wet species (*E. stricta*) to provide broader phylogenetic context. Above the species level, *E. moluccana*, *E. largiflorens*, *E. sideroxylon* and *E. viridis* belong to the Adnataria section, *E. dealbata* and *E. vicina* belong to the Exsertaria section and *E. nitida*, *E. regnans* and *E. stricta* belong to the Eucalyptus section.

For each species, we selected a single site within its natural distribution for sampling (Table S1). For three species pairs (*E. moluccana–E. largiflorens*, *E. dealbata–E. vicina*, and *E. nitida–E. regnans*), we sampled ten individuals per species for DNA extraction (n = 10 per species), enabling population-level analyses of genetic diversity and differentiation. For the remaining species pair (*E. sideroxylon*–*E. viridis*) and *E. stricta*, DNA sampling was limited to one individual per species (Supplementary Table S1). While all nine species provide evolutionary context, the analyses of genome-wide diversity, divergence, and selection presented here focus on the three species pairs with population-level sampling. The species with single individuals are included in complementary genomic analyses in a companion study (James et al., unpublished), which employs methods that do not require within-population sampling.

### DNA extraction, sequencing and bioinformatics

All nine species were sampled within their native distribution during the 2022/2023 austral summer-autumn. For each individual, leaf material for DNA extraction was sampled from young, unexpanded leaves, which were placed inside a small seed envelope, and immediately surrounded with silica beads and stored in an airtight container. For each individual, 20-30 mg of dried leaf material was weighed into a 2 mL microcentrifuge tube. Leaf samples were sent to the Australian Genome Research Facility for DNA extractions and library preparation with the Illumina DNA Prep M Library Prep kit. Samples were pooled in equimolar concentrations and sequenced on two lanes of the NovaSeq 6000 S4, 300 cycle with 150bp paired-end sequencing. On average we obtained 30.8Gb (±7.5 SD) of raw sequencing data per individual.

Raw reads were quality-trimmed using fastp v0.23.4 (S. Chen et al., 2018) with the following parameters: minimum quality score of 10, maximum uncalled bases of 50, and minimum read length of 50bp. Using BWA-MEM v0.7.19 (Li, 2013), quality-trimmed reads were mapped to the *Eucalyptus grandis* v2.0 reference genome (Bartholomé et al., 2015; Myburg et al., 2014), which was downloaded through the Joint Genome Institute Plant Genome Atlas (Sreedasyam et al., 2023). Read groups were added during mapping, and resulting SAM files were converted to BAM format and coordinate-sorted using SAMtools v1.19 (Li et al., 2009). As each individual was sequenced on two lanes, the multiple mapped read files per individual were merged using Picard v3.1.1 (Broad Institute, 2019). Picard was used to clean BAM files and mark PCR duplicates.

Using GATK v4.4.0.0 (McKenna et al., 2010), HaplotypeCaller was run per sample in GVCF mode whilst retaining all variant and invariant sites. Individuals were combined per species using GATK CombineGVCFs. For species with more than one individual, joint genotyping was performed using GATK GenotypeGVCFs whilst retaining both variant and invariant sites. All VCF files were concatenated and merged into a single multi-population VCF using bcftools. Note that missing genotypes with depth (DP) = 0 were corrected into the explicit missing data format (“./.”) using bcftools +setGT. This was because GATK v4.4.0.0 encodes missing genotypes as 0/0 with a DP=0 in the format field for single-sample SNP calling with HaplotypeCaller, which can cause downstream issues.

A multi-step filtering approach was applied using GATK VariantFiltration and VCFtools v0.1.17 (Danecek et al., 2011). Following GATK’s hard filtering recommendations, sites were flagged using GATK VariantFiltration with the following thresholds: quality by depth < 2.0, strand odds ratio > 3.0, Fisher strand bias > 60.0, mapping quality < 30.0, mapping quality rank sum test < -12.5, and read position rank sum test < -8.0. Sites failing these filters were removed using VCFtools. Sites were retained if genotyped in ≥ 50% of individuals and genotypes with < 3 reads were recoded as missing data. Mean depth per site was calculated, and sites with minimum mean depth < 10 and maximum mean depth >100 were removed. Missing data was calculated per species, and sites with > 70% missing data in any species were excluded to ensure representation across all taxa. Finally, sites with > 20% overall missing data and indels were removed, and only biallelic SNPs were retained. We restricted all further analyses to the 11 chromosomes of the *E. grandis* reference genome and removed all scaffolds. The filtering retained 160,686,818 variant and invariant sites across the genome, of which 12,608,766 were variant with a MAF ≥ 0.05.

### Phylogeny

We inferred phylogenies from our dataset to compare to the phylogeny of Thornhill et al., (2019) and to verify that all individuals were correctly assigned to their respective species. We used a multi-step multispecies coalescent approach, in which gene trees were first inferred from multiple sequence alignments, followed by inference of species trees from these gene trees, including all 63 individuals across all nine species.

We used gene coordinates from the *Eucalyptus grandis* v2.0 reference genome to extract VCF files for each of the 34,121 genes on the 11 chromosomes using bcftools. We used the multi-population VCF file with variant and invariant sites for all individuals. VCF files were converted into multiple sequence alignments (MSAs) in PHYLIP format using vcf2phylip v2.0 (Ortiz, 2019), with IUPAC ambiguity codes used for heterozygous sites.

Of the 34,121 gene MSAs, 3,379 contained no data. From the remaining 30,742 MSAs, gene trees were inferred for all individuals using IQ-TREE v2.3.6 (Minh et al., 2020) We used the GTR+Gamma+I substitution model and collapsed near-zero branches using the - czb option. We used the default minimum branch length of the smaller of 0.000001 and 0.1/alignment_length. With the longest alignment length being 49,002 sites, the minimum branch length defaulted to 0.000001 for all genes. For further control of the gene tree topologies, we used the collapseEdges function of the R package MSCquartets (Rhodes et al., 2021) to collapse branches with lengths less than 0.000001 to polytomies.

We filtered gene trees based on alignment length and taxon occupancy. Previous studies have shown that filtering based on taxon occupancy has variable effects on species tree inference accuracy and can be neutral or detrimental (Hosner et al., 2016; Jiang et al., 2014; Molloy & Warnow, 2018; Streicher et al., 2016). In contrast, filtering gene trees based on gene tree estimation error, often associated with shorter alignments and reduced phylogenetic signal, is generally beneficial (Blom et al., 2017; M.-Y. Chen et al., 2015; Hosner et al., 2016; Simmons et al., 2016). However, Molloy & Warnow (2018) showed that the effects of such filtering are context dependent and may be beneficial or detrimental depending on the level of incomplete lineage sorting, with greater benefits expected in low incomplete lineage sorting scenarios. We therefore evaluated five filtering scenarios spanning a gradient of stringency: 1) no filtering (30,742 genes), 2) at least 100 bp and at least 51 individuals (29,288 genes), 3) at least 250 bp and at least 57 individuals (26,950 genes), 4) at least 500 bp and at least 60 individuals (23,377 genes), and 5) at least 1000 bp and all 63 individuals (18,658 genes).

Species trees were inferred for each set of gene trees using ASTRAL-IV (Tabatabaee et al., 2023; Zhang et al., 2025). ASTRAL-IV is a software based on the multispecies coalescent model of incomplete lineage sorting. It infers an unrooted species tree from a set of gene trees. On the inferred species trees, we did not enforce an outgroup or constrain individuals of the same species to form clades, in order to independently verify species assignments.

### Population structure

To further explore clustering of populations, we undertook a Principal Components Analysis (PCA) on SNPs filtered for a MAF ≥ 0.05. SNPs were pruned for linkage disequilibrium in PLINK v1.9 (Purcell et al., 2007) using a sliding window of 50 SNPs, advanced in steps of 10 SNPs, retaining SNPs with pairwise r² < 0.1. This resulted in 428,330 SNPs. The PCA was then performed on the resulting pruned dataset in PLINK v1.9, and the variance explained was extracted for the first 20 PCs.

### Genome-wide landscapes of diversity and divergence

To explore diversity and divergence along the genome, we analysed genome-wide patterns of F_ST_ and *d*_XY_ between species, as well as π and Tajima’s D within species. This was undertaken for the three species pairs with 10 individuals per species: the two wet-dry contrasts of *E. moluccana–E. largiflorens* and *E. dealbata–E. vicina*, and the wetter-wet contrast of *E. nitida–E. regnans*. F_ST_, *d*_XY_ and π were calculated with 20kB non-overlapping sliding windows using pixy v1.2.10 (Korunes & Samuk, 2021), which accounts for monomorphic sites when calculating diversity and divergence statistics. Tajima’s D was calculated using VCFtools using the same 20kB non-overlapping sliding windows. To measure concordance across diversity and divergence landscapes, we calculated Pearson’s *r* and Spearman’s *ρ* correlations across the non-overlapping 20kB windows for all diversity and divergence statistics. For visualisation, we smoothed the *d*_XY_, π and Tajima’s D plots by aggregating windows into 1.5Mb bins and calculating the mean statistic per bin. All statistical analyses and data visualisation were undertaken in R v4.1.0 (R Core Team, 2017).

### Linked selection throughout the genome

We calculated population genetics statistics (π, F_ST_ and *d*_XY_) in non-overlapping 1Mb windows across the genome for each species pair (results were consistent when all analysis was repeated for 20kB windows; data not shown). In addition, we calculated (in non-overlapping 1Mb windows): GC content from the *Eucalyptus grandis* v2.0 reference genome using bedtools v2.29.1 (Quinlan & Hall, 2010), gene density from the reference genome using a custom R script, where a gene was considered to reside in a window if more than 50% of its length overlapped the window, and recombination rate variation from the linkage map used to construct the *E. grandis* reference genome (Version 2 markers, GRA parent) (Bartholomé et al., 2015).

To explore the prevalence of linked selection along the genome, we measured correlations between: F_ST_ vs mean π per pair (negative association expected under linked selection as low diversity inflates F_ST_), *d*_XY_ vs mean π per pair (positive association expected under background selection, which depresses both polymorphism and absolute divergence), mean π and gene density (where a negative association is expected because selection at linked sites reduces diversity in gene-dense regions), and mean π and recombination rate (cM/Mb) (where a positive association is expected because recombination reduces the genomic footprint of selection at linked sites). Additionally, we assessed associations between genome features including recombination rates, gene density and GC content. Because recombination rate is itself correlated with both gene density and GC content, we residualised recombination rate (natural log) against gene density and GC content using a generalised additive model with thin-plate smooth terms for each predictor using the mgcv package in R (Wood, 2025), to account for their non-linear relationships with recombination rate. The resulting residuals represent recombination rate variation independent of gene density and GC content, and were used to assess the relationship with mean π, F_ST_ and *d*_XY_. All correlations were visualised as scatterplots, with linear regressions for visualisation. We calculated Pearson’s *r* and Spearman’s *ρ* correlations across the non-overlapping 1Mb windows for all scatterplots.

### Detection of genomic outliers

To examine the repeatability of the genetic basis of drought adaptation, we detected genomic outliers for each species pair, by focussing on residual differentiation after accounting for genome-wide variation in nucleotide diversity. Because F_ST_ is a relative measure of differentiation that can be inflated in regions of low nucleotide diversity due to linked selection, we controlled for this effect by regressing *d*_XY_ on mean π using a linear model and F_ST_ on mean π using LOESS smoothing (span = 0.6), and then extracting the residuals for each window. To confirm that these regressions removed the correlation between differentiation metrics and nucleotide diversity, we calculated Pearson’s *r* between the residuals and mean π. For all three species pairs, *d*_XY_ residuals were uncorrelated with mean π (Pearson’s *r* = 0.00, P-value = 1.00), and F_ST_ residuals showed negligible correlation with mean π (*E. moluccana–E. largiflorens* Pearson’s *r* = 0.01, P-value = 0.05; *E. dealbata–E. vicina* Pearson’s *r* = 0.006, P-value = 0.32; *E. nitida*–*E. regnans* Pearson’s r = 0.02, P-value = 0.20). Although *d*_XY_ does not mathematically depend on π, both metrics are strongly correlated because local effective population size (Ne) and linked selection jointly influence polymorphism and divergence; adjusting for π removes this confounding effect.

Outlier windows were defined as those falling within the top 1% for both residual F_ST_ and residual *d*_XY_, a conservative threshold that represents regions where relative and absolute divergence are jointly elevated beyond expectations from background selection. To validate our residual-based approach, we also undertook additional outlier analysis where, for each species pair, we considered a window an outlier if it fell within the top 10% of F_ST_ windows as well as the top 10% of *d*_XY_ windows (without controlling for the residuals). This threshold of 10% ensured a sufficient number of outlier windows were detected, as more stringent thresholds yielded very few windows with jointly elevated F_ST_ and *d*_XY_. A gene was considered to be located within a genomic window if at least 25% of its length resided within the window.

### Gene ontology enrichment analysis

To determine whether outlier genes were enriched for any biological functions, we performed gene ontology (GO) enrichment analysis using the web-based version of DAVID v6.8 (D. W. Huang et al., 2009a, 2009b). We undertook this analysis for each of the three species pairs. For each outlier gene set, we obtained the *Arabidopsis* orthologues as outlined above and used these orthologues as the target gene set, and all *Arabidopsis* orthologues present in the *E. grandis* reference genome (one orthologue per gene) as the background gene set. A GO term was considered enriched if it had a P-value < 0.05 (the EASE score, calculated using a modification of Fisher’s exact test; Huang et al., 2009a, 2009b). We assessed enrichment at the level of the biological processes (*GOTERM_BP_DIRECT*).

### Association of outliers with genome features

For each species pair, we tested whether outlier windows were associated with a range of genome features, including nucleotide diversity (π) in the wet and dry populations, recombination rate, gene density, GC content, F_ST_ and *d*_XY_, calculated as described above. For each genome feature, we compared outlier windows against the remaining (background) windows using a permutation test. We took the median of the outlier windows as the test statistic and generated a null distribution by drawing 10,000 random samples of background windows, each matched to the number of outlier windows, and recording the median of each sample. A two-sided empirical P-value was calculated as the proportion of null medians at least as extreme as the observed median. This analysis was performed separately for the two outlier sets (windows in the top 1% for both residual F_ST_ and residual *d*_XY_, and windows in the top 10% for both F_ST_ and *d*_XY_).

### Genomic islands of differentiation

To complement the window-based outlier analysis, we characterised broad genomic islands of differentiation, defined as extended contiguous regions of elevated F_ST_ rather than individual outlier windows. Such islands can arise through divergent selection, but also through linked selection acting on regions of reduced recombination, and examining patterns of diversity and linkage disequilibrium within them can help characterise their underlying structure. Genomic islands were defined as regions in the top 5% of the genome-wide F_ST_ distribution spanning at least ten consecutive 20kb windows, allowing for up to two consecutive non-outlier windows within each island. Windows lacking data were excluded from the analysis. For each species pair, we performed GO enrichment analysis of the genes residing within the islands, as described above, and focused subsequent analyses on the three largest islands. For these islands, patterns of linkage disequilibrium (LD) were examined across each island and its symmetric flanking regions, extending a distance equal to 60% of the island length on either side. SNPs were thinned in PLINK to retain one SNP every 5kb, and pairwise LD was calculated as r² between all pairs of thinned SNPs, then summarised for visualisation into 10 kb bins by calculating the mean r² per bin. Population differentiation (F_ST_) and nucleotide diversity (π) were visualised across each island and its symmetric flanking regions, with values smoothed using a rolling mean across 25 consecutive windows.

Finally, we assessed whether islands were associated with particular features of the genomic landscape. We examined whether islands preferentially occurred in regions of reduced recombination and elevated gene density, as expected if linked selection contributes to their formation, and whether islands showed the reduced π and comparable *d*_XY_ expected under linked selection, or the elevated *d*_XY_ expected under increased divergence. We compared each island against the genome background. Background windows were defined as all windows not falling within any island. For each island and each genomic feature, we tested whether the distribution of values across the island’s windows differed from that of the background windows using a two-sided Wilcoxon rank-sum test. To account for multiple testing across islands, P-values were adjusted within each feature and species pair using the Benjamini–Hochberg false discovery rate.

## RESULTS

### Phylogenetic relationships and population structure

Both the inferred phylogenies and principal components analysis supported the independent origin of the study lineages: each of the three species pairs belongs to a distinct taxonomic section (Adnataria, Exsertaria, and Eucalyptus), with species grouping by section rather than by the ecological habitat in which they are found (Figure 1; Figure S1).

The inferred species trees were highly congruent across filtering scenarios, differing primarily in levels of branch support and placement of individuals within species clades. Overall, the phylogenies recovered consistent relationships among species and strong support for most clades. Across all five species trees, individuals from each multi-sample species formed monophyletic groups. When considering any one individual per species, all five species trees were topologically identical. For visualisation purposes, we used the prior knowledge of the *Eucalyptus* section (*E. nitida, E. regnans and E. stricta*) being an outgroup to root the phylogenies. Almost all clades were very well supported, with local posterior probabilities of 1. The exception is the *E. moluccana* and *E. largiflorens* clade, where the local posterior probabilities range from 0.00434 (scenario 5) to 0.909 (scenario 2). This is to be expected, as filtering genes based on missing data can decrease branch support in ASTRAL (Molloy & Warnow, 2018).

The inferred phylogenies were largely congruent with the phylogeny of Thornhill et al., (2019), with the exception of the placement of *E. viridis* and *E. sideroxylon*. Thornhill et al., (2019) placed these species as sister taxa within the Adnataria section. Our phylogenies inferred *E. sideroxylon* as diverging first, followed by *E. viridis* and finally by *E. moluccana* and *E. largiflorens*. We had only one individual each for *E. viridis* and *E. sideroxylon*, which could have impacted the accuracy of the placement of these species on the phylogeny.

### Genome-wide diversity and divergence landscapes

Genome-wide scans revealed heterogeneous landscapes of diversity and differentiation across all three species pairs (Figure S2). Average nucleotide diversity (π ± SE) was similar across species (*E. moluccana* 0.022 ± 0.00008; *E. largiflorens* 0.024 ± 0.00008; *E. dealbata* 0.018 ± 0.00007; *E. vicina* 0.018 ± 0.00007; *E. nitida* 0.021 ± 0.00008, *E. regnans* 0.024 ± 0.00008). There were very strong correlations of π landscapes across all species (mean Pearson’s *r* = 0.70, range = 0.51-0.89; mean Spearman’s *ρ* = 0.77, range 0.69-0.91), and weak to moderate correlations of Tajima’s D (mean Pearson’s *r* = 0.21, range = 0.10-0.45; mean Spearman’s *ρ* = 0.21, range 0.10-0.45) (Table S2). *d*_XY_ landscapes between pairs were strongly correlated (Pearson’s *r* range = 0.64-0.75, Spearman’s *ρ* = 0.70-0.79), and F_ST_ landscapes were weakly to moderately correlated (Pearson’s *r* = 0.18-0.50, Spearman’s *ρ* = 0.23-0.56) (Table S2). Together, the strong correlations of both π and *d*_XY_ landscapes suggests that similar evolutionary processes, particularly linked selection (where diversity is reduced in low-recombination regions), shape genome-wide landscapes of diversity across the *Eucalyptus* lineages.

### Pervasive linked selection throughout the genome

To further explore the prevalence of linked selection along the genome, we assessed correlations between population genetics statistics and genome features (Figure 2, Figure S3, and see Table S2 for all statistics). F_ST_ was negatively correlated with mean π in all pairs (Figure 2A), indicating that elevated relative differentiation tends to occur in regions of reduced absolute diversity, consistent with low-diversity regions inflating F_ST_. Consistent with linked selection reducing genome-wide diversity, we observed very strong positive correlations between *d*_XY_ and mean π across all three species pairs (Figure 2B), indicating that regions of reduced diversity also show reduced absolute divergence. Mean π was negatively correlated with gene density across all pairs (Figure 2C), consistent with selection at linked sites reducing diversity in gene-dense regions. Mean π was also negatively correlated with GC content across all pairs (Figure S4A), consistent with reduced diversity in gene-dense regions, as GC content is itself positively correlated with gene density in Eucalyptus (Figure S5). However, as gene density and GC content are themselves positively correlated with recombination rate (Figure S5). However, this bivariate correlation between gene density and GC content should be interpreted with caution, as both features are themselves correlated with recombination rate.

**Figure 2.**
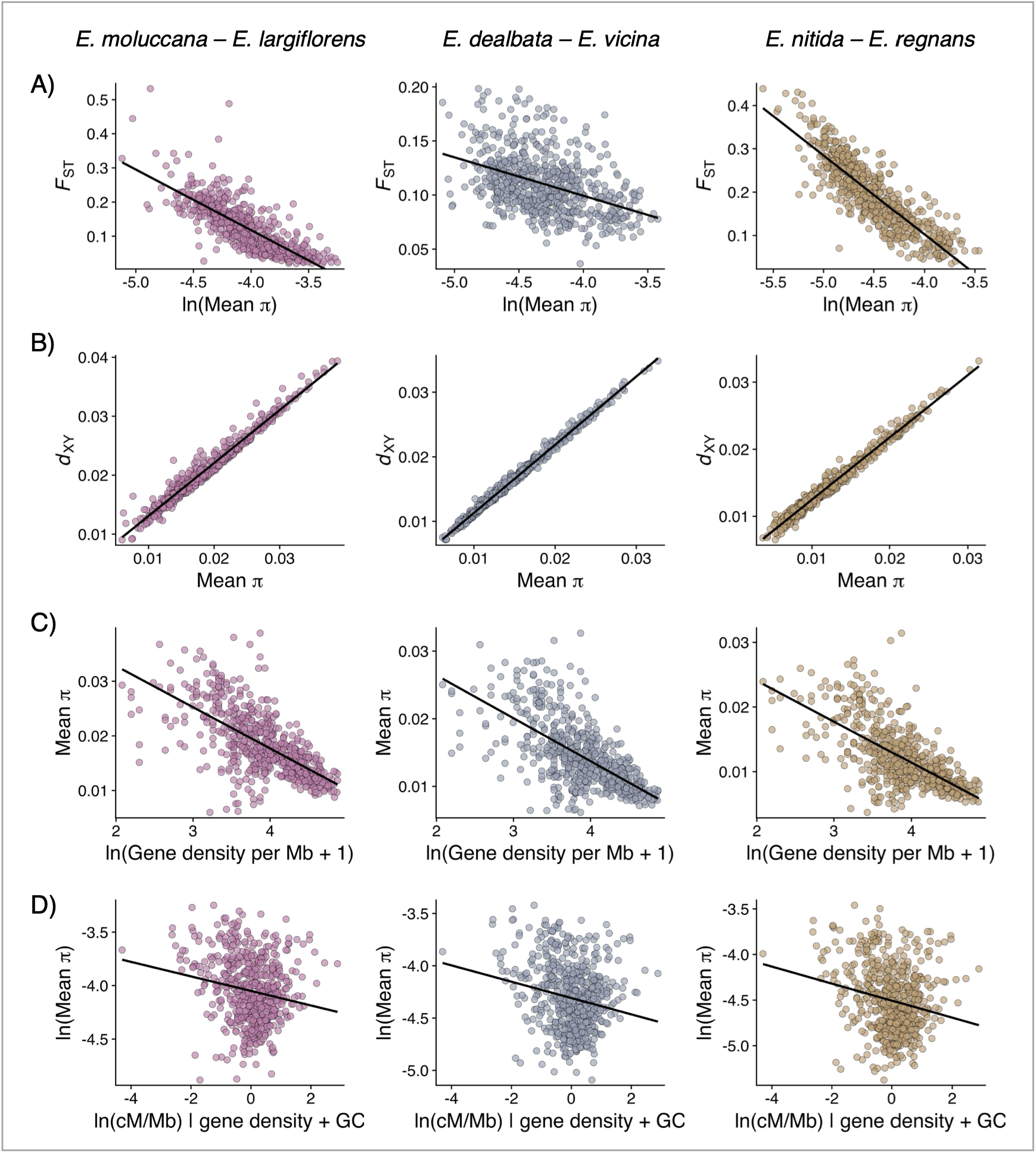
Evidence for linked selection. Scatterplots of pairwise relationships across the genome for *E. moluccana*–*E. largiflorens* (pink), *E. dealbata*–*E. vicina* (slate) and *E. nitida*–*E. regnans* (brown) for A) F_ST_ vs mean π, B) *d*_XY_ vs mean π, C) Mean π vs gene density per Mb, and D) mean π vs recombination rate, after accounting for gene density and GC content. Each point represents a 1Mb non-overlapping genomic window; trend lines show linear regressions for visualisation. See Table S2 for Pearson’s and Spearman’s correlation coefficients and associated P-values.

We next examined how recombination rate relates to diversity and divergence, both directly and after accounting for gene density and GC content (Figure 2D, Figure S6, Table S2). Contrary to the simple expectation under linked selection, where higher recombination should reduce the genomic footprint of selection and therefore elevate diversity, mean π was negatively correlated with recombination rate across all three species pairs (Figure S4B). Absolute divergence (*d*_XY_) was likewise negatively correlated with recombination rate (Figure S6A), whereas F_ST_ was positively correlated with recombination rate (Figure S6B). These relationships were largely retained after conditioning recombination rate on gene density and GC content: mean π (Figure 2D) and *d*_XY_ (Figure S6C) remained negatively associated with recombination, and F_ST_ (Figure S6D) remained positively associated, indicating that these patterns are not simply a by-product of the collinearity between recombination, gene density and GC content.

### Non-convergent signatures of selection across species pairs

We next identified genomic outliers for each species pair by examining windows in the top 1% for both F_ST_ and *d*_XY_, after accounting for genome-wide variation in nucleotide diversity (see Methods). Because F_ST_ can be inflated in regions of low diversity, raw F_ST_ and *d*_XY_ need not coincide, and elevated relative differentiation does not necessarily reflect elevated absolute divergence. Controlling for π should therefore reveal whether regions of high differentiation also show elevated divergence beyond the effects of linked selection. Consistent with this, after accounting for variation in π through residual analysis, F_ST_ and *d*_XY_ residuals showed strong positive correlations for all three species pairs (Figure S7), suggesting that once background diversity levels are controlled for, regions with elevated absolute divergence also show elevated relative differentiation. This supports the use of joint F_ST_ and *d*_XY_ residual outliers to identify candidate regions of divergence.

The number of outlier windows was similar across species pairs (Figure 3A-C): 159 outlier windows for *E. moluccana*–*E. largiflorens* (containing 119 genes, 99 with *Arabidopsis* orthologues; Table S3), 155 for *E. dealbata*–*E. vicina* (containing 91 genes, 78 with *Arabidopsis* orthologues; Table S4), and 161 for *E. nitida–E. regnans* (containing 155 genes, 137 with *Arabidopsis* orthologues; Table S5). The outlier windows were distributed across all chromosomes, with visually notable clustering on chromosomes two and three for the *E. moluccana*–*E. largiflorens* species pair (Figure 3A-C).

**Figure 3.**
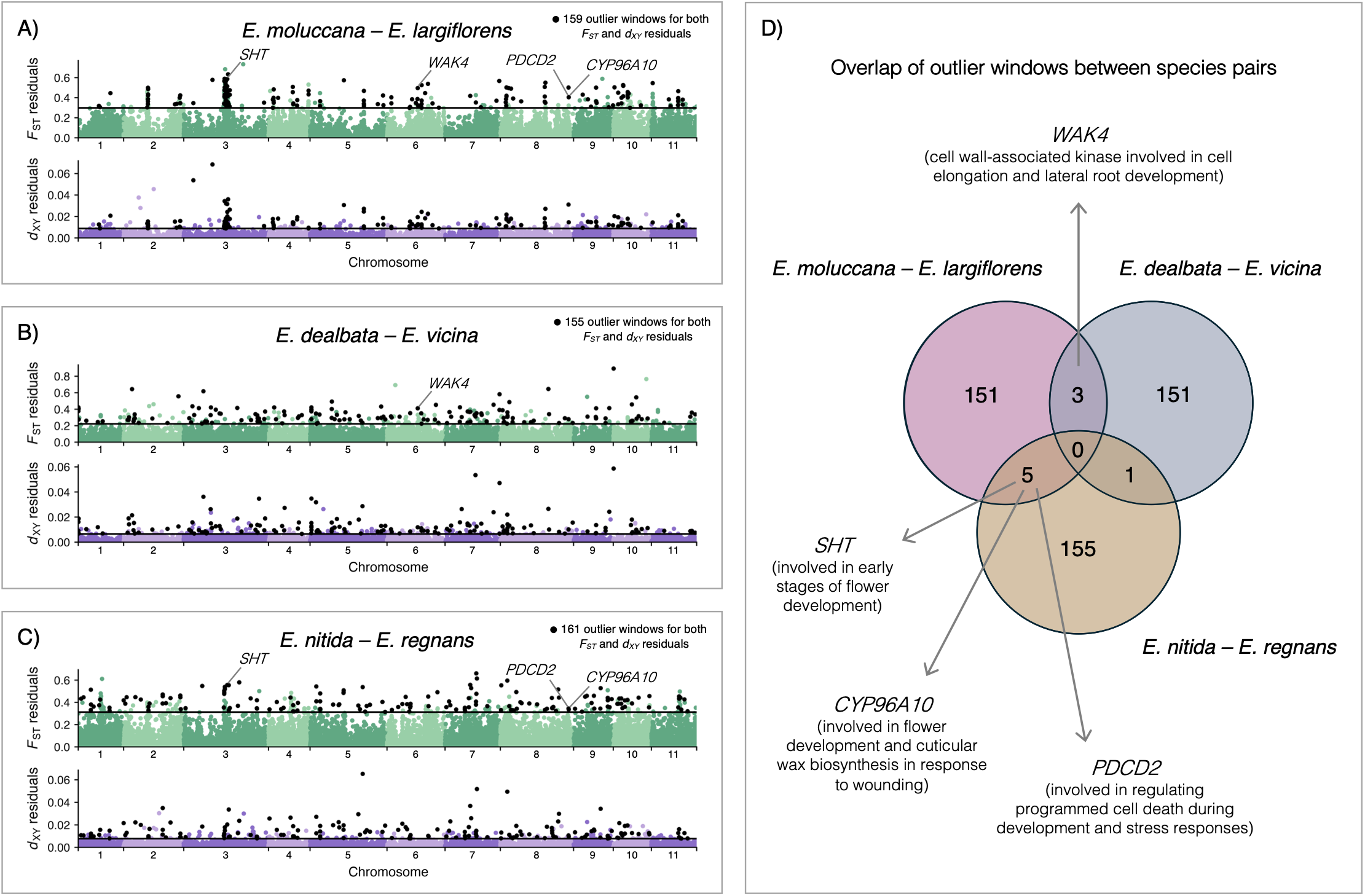
Shared signatures of selection across three replicate species pairs. F_ST_ and *d*_XY_ residuals after c *Eucalyptus* chromosomes for A) *E. moluccana*–*E. largiflorens*, B) *E. dealbata*–*E. vicina* and C) *E. nitida*–*E.* indicate the significance threshold (top 1% of genomic windows). Each dot represents a 20kB non-overlappi represent outlier windows found in the top 1% of both F_ST_ and *d*_XY_ residuals. For ease of visualisation of the zero. D) Overlap of outlier windows across the three species pairs. Arrows and gene descriptions represent g windows.

Surprisingly, we detected very few shared outlier windows between pairs (Figure 3D). This pattern suggests that while each drier-adapted species faces similar environmental challenges, they have evolved distinct genetic solutions to aridity rather than repeatedly utilising the same genomic regions. Only three outlier windows were shared between *E. moluccana*–*E. largiflorens* and *E. dealbata*–*E. vicina*, with just one containing a gene: *WAK4* (a cell wall-associated kinase involved in cell elongation and lateral root development). Five outlier windows were shared between *E. moluccana*–*E. largiflorens* and *E. nitida–E. regnans*, harbouring genes including *SHT* (involved in early stages of flower development), *PDCD2* a programmed cell death 2 C-terminal domain-containing protein (involved in regulating programmed cell death during development and stress responses), and *CYP96A10* (involved in flower development and cuticular wax biosynthesis in response to wounding). One outlier window was shared between *E. dealbata*–*E. vicina* and *E. nitida–E. regnans*, although this window contained no genes. No outlier windows were shared across all three pairs. Together, this limited overlap suggests that independent transitions into drier environments have been achieved through largely distinct genetic routes, indicating that the genetic basis of drought adaptation in *Eucalyptus* is not strongly constrained to specific genomic regions. The wetter-wet pair (*E. nitida*–*E. regnans*), which does not involve adaptation to semi-arid conditions, also showed minimal overlap with the wet-dry pairs, suggesting that genomic divergence along precipitation gradients (whether involving aridity or not) tends to involve lineage-specific genomic regions.

### Some convergence of outlier biological functions

Gene ontology enrichment analysis of genes within outlier windows revealed partially overlapping functional profiles for each species pair (Table S6). The *E. moluccana*–*E. largiflorens* pair was enriched for functions related to metabolic processes, including gibberellin and fatty acid metabolism. The *E. dealbata*–*E. vicina* pair showed enrichment for biotic stress responses, including defence response and response to oomycetes and other organisms. The *E. nitida*–*E. regnans* pair was enriched for defence-related hormone and secondary metabolite processes, including defence response, and lipid, flavonoid, and jasmonic acid biosynthesis. Notably, only one biological process was significantly enriched across multiple pairs: defence response, which was enriched in both the *E. dealbata*–*E. vicina* and *E. nitida*–*E. regnans* comparisons.

To validate our residual-based outlier approach, we compared it with a traditional method where outlier windows were defined as those in both the top 10% of raw F_ST_ and *d*_XY_ values. This approach again identified outliers distributed across all chromosomes, with minimal overlap of outlier windows (Figure S8, Tables S7-9). Gene ontology enrichment analysis revealed largely concordant functional profiles between the two methods: each pair showed enrichment for similar biological processes as observed with the residual approach, and defence response was again the only function shared between the *E. dealbata*–*E. vicina* and *E. nitida*–*E. regnans* pairs (Table S10).

### Genomic features of outlier regions

For each species pair, we compared outlier windows (in the top 1% for both residual F_ST_ and *d*_XY_) against the genomic background across a range of genome features (Figure 4, Figure S12). Outlier windows were characterised by significantly reduced nucleotide diversity in both the wet and dry populations of *E. moluccana*–*E. largiflorens* and *E. dealbata*–*E. vicina*, yet outliers in the *E. nitida*–*E. regnans* pair contained elevated nucleotide diversity (Figure 4A,B). Recombination rate was significantly reduced in outlier windows, for only the *E. nitida*–*E. regnans* pair (Figure 4C). Gene density was significantly reduced in outliers of *E. moluccana*–*E. largiflorens* and *E. dealbata*–*E. vicina* (Figure 4D). GC content was significantly reduced in outliers for the *E. dealbata*–*E. vicina* and *E. nitida*–*E. regnans* pairs (Figure S12C). Similar patterns were recovered using the top 10% outlier set (Figures S13, S14).

**Figure 4.**
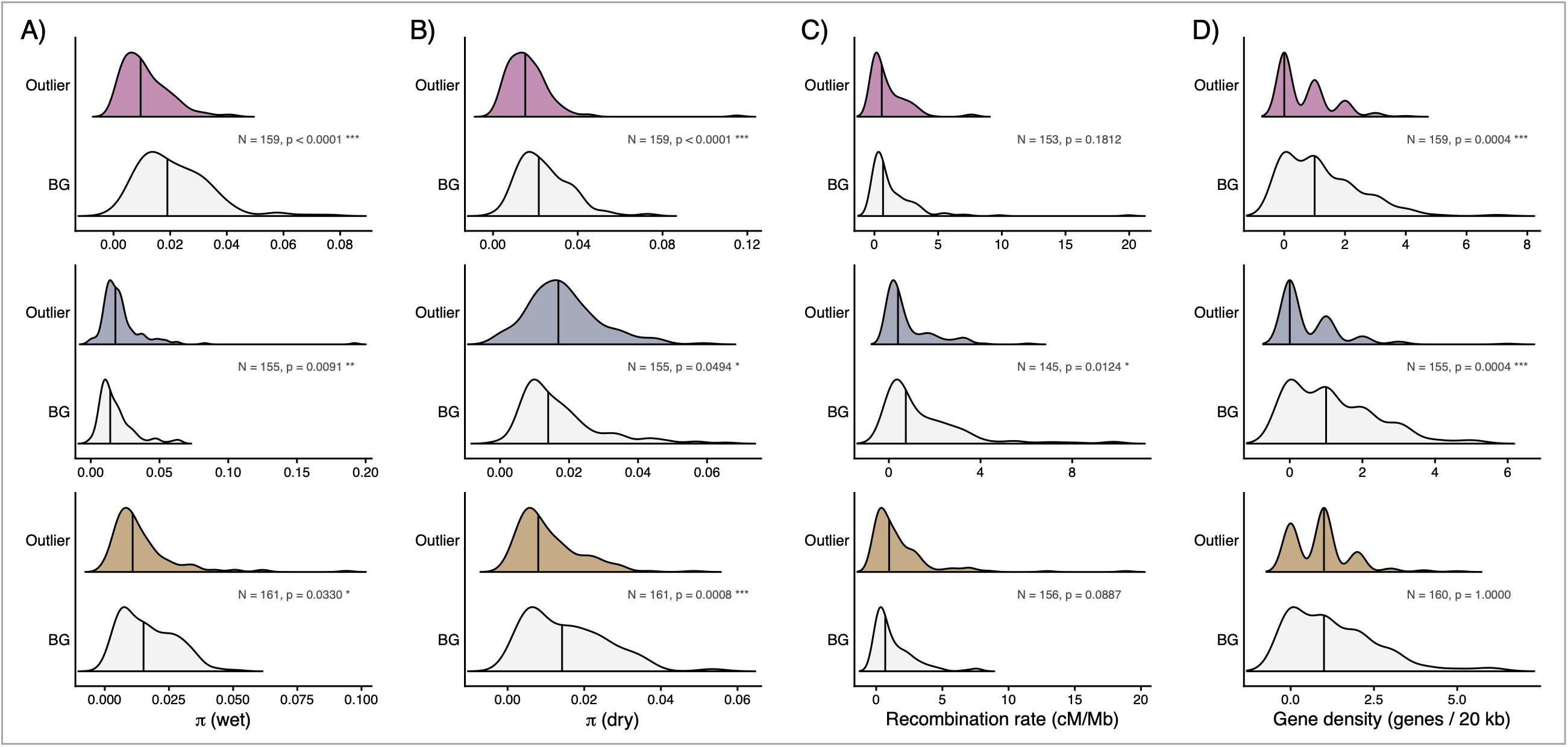
Associations of outliers with genome features. Half-violin plots for outlier windows (top, colour background windows (BG, bottom, grey) for A) nucleotide diversity (π) in the wet-adapted species, B) π in t recombination rate (cM/Mb), and D) gene density (genes per 20 kb). Vertical bars represent the median of ea permutation test comparing outlier and background windows, in which the observed median of the outlier wi distribution generated by drawing 10,000 random samples of background windows matched to the number of outlier windows and two-sided empirical P-value; * p < 0.05, ** p < 0.01, *** p < 0.001). Top row: *E. moluc* row: *E. dealbata*–*E. vicina* (slate); bottom row: *E. nitida*–*E. regnans* (brown). Outlier windows were defined F_ST_ and *d*_XY_ residuals.

### Large genomic islands of differentiation

We examined genomic islands of differentiation by identifying clusters of elevated F_ST_ along the genome. Some clusters occurred in close succession, separated by gaps of only three to four genomic windows, and formed what visually appeared as a single block in the F_ST_ profiles. We therefore merged each such pair into a single island. In total we identified 23 islands across the three species pairs, ranging from 280kb to 2.24Mb in length (Table S11); with 11 islands in *E. moluccana*–*E. largiflorens*, 10 in *E. nitida*–*E. regnans*, and 2 in *E. dealbata*–*E. vicina*. To assess whether these islands coincided with features of the genomic landscape, we compared each island against the genomic background for nucleotide diversity (π) in each population, *d*_XY_, recombination rate, gene density and GC content, testing each island’s windows against all background windows (Figure 5A).

**Figure 5.**
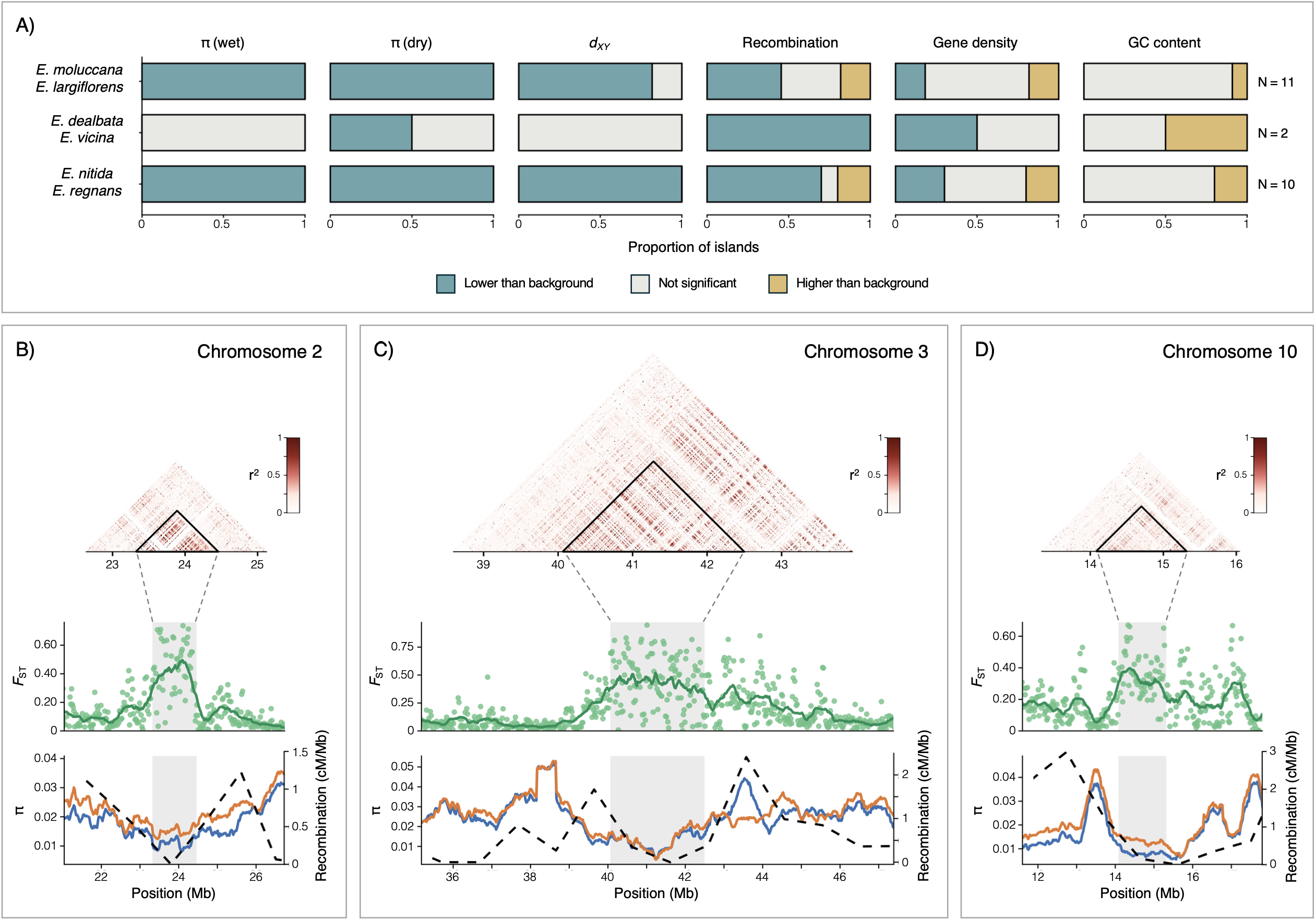
Islands of divergence and their associated genome features. A) Associations of islands of diver three species pairs (*E. moluccana*–*E. largiflorens*, *E. dealbata*–*E. vicina* and *E. nitida*–*E. regnans*). For each the distribution of values across the island’s windows was compared with that of the background windows (a island) using a two-sided Wilcoxon rank-sum test; P-values were adjusted within each feature and species pa false discovery rate. Bars show the proportion of islands that were significantly lower than background (teal) background (grey), or significantly higher than background (mustard). B-D) Patterns of linkage disequilibriu (F_ST_), nucleotide diversity (π) and recombination for the three largest islands of divergence and flaking regio *moluccana*–*E. largiflorens* species pair on: B) chromosome two, C) chromosome three, and D) chromosome black triangle (LD plot), or with grey shading (F_ST_, π and recombination plots). Each dot represents a 20kB n recombination values were smoothed using a rolling mean across 25 consecutive windows and connecting ad wetter species is in orange, π for the drier species is blue).

We observed that π was significantly reduced within islands relative to background in every island in both the wet and dry populations of *E. moluccana*–*E. largiflorens* (11 of 11) and *E. nitida*–*E. regnans* (10 of 10). *d*_XY_ was likewise reduced within most islands (9 of 11 in *E. moluccana*–*E. largiflorens* and 10 of 10 in *E. nitida*–*E. regnans*, with the remainder not differing significantly), and no island showed elevated *d*_XY_. This combination of reduced π together with reduced or comparable *d*_XY_ is consistent with linked selection reducing diversity within islands, rather than with elevated between-lineage divergence. Recombination rate was significantly reduced within islands for 5 of 11 islands in *E. moluccana*–*E. largiflorens*, 2 of 2 in *E. dealbata*–*E. vicina*, and 7 of 10 in *E. nitida*–*E. regnans* (with a few islands showing elevated recombination in the *E. moluccana*–*E. largiflorens* and *E. nitida*–*E. regnans* pairs). Gene density and GC content showed no consistent association with islands, where most islands did not differ significantly from the background.

The three largest islands resided in the *E. moluccana*–*E. largiflorens* pair on chromosome two (Figure 5B), three (Figure 5C) and ten, (Figure 5D). In each island, elevated F_ST_ across the island coincided with an increase in linkage disequilibrium, visible as a dense block of high pairwise r² spanning the island in the LD heatmaps. Within each island, nucleotide diversity (π) was noticeably reduced in both populations relative to the surrounding regions, and recombination rate was correspondingly low across the island before rising in the flanking regions. This combination of reduced diversity and low recombination within regions of high differentiation is consistent with a role for linked selection in shaping the largest islands.

Gene ontology enrichment analysis of island genes revealed pair-specific functional enrichment (Table S12). In *E. moluccana–E. largiflorens*, the most enriched processes included the regulation of transcription, leaf senescence, response to jasmonic acid, and primary cell wall biogenesis, alongside several stress-related categories (cellular responses to hypoxia and freezing). The *E. nitida–E. regnans* islands were enriched for regulation of transcription, callus formation, root development and starch biosynthesis, while *E. dealbata–E. vicina*, which contained only two islands, showed a single weakly enriched category (polysaccharide catabolism). No process was significantly enriched across all three pairs, consistent with the largely lineage-specific islands.

Individual islands harboured genes with well-characterised roles in abiotic-stress and drought responses (see Table S13). Notably, several of these were transcription factors central to the drought and abscisic acid (ABA) regulatory network. Islands in *E. moluccana–E. largiflorens* contained: *MYC2*, a bHLH transcription factor induced by dehydration and ABA that activates drought-responsive genes (Kazan & Manners, 2013), *VNI2* an ABA-responsive NAC factor that integrates abiotic-stress signals into leaf senescence via the *COR*/*RD* dehydration genes (S.-D. Yang et al., 2011), and *DRIP1*, a RING E3 ligase that regulates the drought master-regulator DREB2A (Qin et al., 2008). In *E. nitida–E. regnans*, an island contained *CBF4*, a C-repeat binding factor that is specifically up-regulated by drought and whose overexpression confers drought tolerance (Haake et al., 2002).

## DISCUSSION

Understanding how independent lineages respond to shared environmental challenges is central to determining whether the genetic basis of adaptation is repeatable. Here we characterised the genomic landscape of divergence across three independent *Eucalyptus* species pairs that transitioned into drier environments. We found pervasive signatures of linked selection, which shaped diversity and differentiation across all pairs and produced broadly concordant genomic landscapes, as well as large regions of genomic differentiation. Although these shared genomic architectures have the potential to create the appearance of convergent adaptation due to differentiation occurring in the same genomic regions across lineages, our candidate regions associated with the transition to drier environments were largely non-convergent across pairs, with little repeatability of selection upon individual loci. We instead observed some convergence at the level of the biological function. Below, we discuss these results and their implications for our understanding of the genomic basis of adaptation.

### Pervasive linked selection throughout the genome

Across all three species pairs we observed genomic signatures consistent with pervasive linked selection. The strong concordance of nucleotide diversity (π) landscapes and absolute divergence (*d*_XY_) landscapes among species implies long-term and persistent linked selection acting on shared features of the genome before the species diverged. These strong correlations of landscape features are similar to patterns observed across the divergence continuum in *Populus* plants (Shang et al., 2023), *Castanopsis* trees (X.-Y. Chen et al., 2024) and stonechat birds (Van Doren et al., 2017). Additionally, we observed that diversity was reduced in regions of high gene density and was positively correlated with *d*_XY_, both hallmarks of selection at linked sites removing variation in and around functional regions (Begun & Aquadro, 1992; Charlesworth et al., 1993; Cutter & Payseur, 2013). The positive correlation of *d*_XY_ and π also implies background selection as a dominant force shaping genome-wide diversity, since the reduction of diversity in the same regions before and after divergence is expected under purifying selection against deleterious mutations rather than selective sweeps (Charlesworth et al., 1993; Cutter & Payseur, 2013). Our observed negative correlations between F_ST_ and π further indicate that elevated relative differentiation tends to arise in low-diversity regions, consistent with the inflation of F_ST_ where within-population diversity is low (Cruickshank & Hahn, 2014). Together, these patterns provide strong evidence that linked selection acting in low-recombination regions is a major determinant of diversity levels across the *Eucalyptus* genome, and that the broad architecture of the differentiation landscape is shaped by shared genomic features.

However, we observed that recombination rate does not relate to diversity and divergence in the manner predicted by the simple model of linked selection, in which recombination is expected to be positively associated with diversity and negatively associated with relative differentiation. Instead, in *Eucalyptus*, regions of high recombination were characterised by reduced diversity and elevated F_ST_. Crucially, this pattern persisted after controlling for gene density and GC content across all three species pairs, indicating that it is not simply a by-product of collinearity among these features. One possibility is that because high-recombination regions in *Eucalyptus* coincide with a high density of targets for selection, the intensity of linked selection does not decline with recombination in the expected way. This interpretation is supported by previous work in *Eucalyptus*, where a negative correlation between diversity and recombination was reported across the genus and attributed to the tight coupling of recombination with gene density and GC content (Gion et al., 2016). Because recombination is so tightly bound to gene density in *Eucalyptus*, high-recombination regions are also high-selection-target regions, which may flip the usual correlation between diversity and recombination. Furthermore, the relationship between recombination and diversity may be more complex, as recombination rate is itself shaped by transposable-element density (Y. Huang et al., 2025) and can be directly mutagenic, contributing to the covariation between recombination, diversity, and divergence (Halldorsson et al., 2019; Kulathinal et al., 2008).

An alternative explanation is that our reliance on a recombination map derived from the *E. grandis* reference genome does not fully capture the recombination landscape of each species pair. Recombination rates themselves are highly dynamic and are known to vary substantially among species and across genomic regions (Ortiz-Barrientos & James, 2017; Stapley et al., 2017). However, in *Eucalyptus* species the relative recombination rate of each chromosome appears highly conserved (Gion et al., 2016), suggesting that a reference-derived map is reasonably transferable at the chromosomal scale. Yet even when chromosome-scale recombination rates are conserved, fine-scale recombination can substantially diverge between lineages, as seen in flycatchers (Kawakami et al., 2017). As a result, our chromosome-scale recombination map may not resolve the fine-scale variation in recombination that locally shapes linked selection, and the local relationship between recombination and diversity within each pair may be obscured. Nevertheless, the strong concordance of genome-wide patterns of nucleotide diversity and absolute divergence across species pairs suggests that the recombination landscape and linked selection exert a consistent influence on patterns of genomic variation.

### Multiple solutions to the same problem

As linked selection acts on shared genomic features, it can drive independent lineages towards similar differentiation landscapes, which can create the appearance of convergent adaptation (Renaut et al., 2013). Despite this shared genomic background, the regions most strongly associated with the transition to drier environments were remarkably non-convergent across pairs: outlier windows were distributed across all chromosomes in each pair, yet we observed a near-absence of shared outliers, indicating that independent transitions to drier environments have occurred through largely distinct genetic routes. Such a pattern is consistent with a highly polygenic architecture of climate adaptation in *Eucalyptus*, in which drought response is coordinated across many genes of small effect scattered throughout the genome, as detected in previous work (Ammitzboll et al., 2020; Butler et al., 2022; Steane et al., 2017). Although previous work shows that drought-resilience strategies in *Eucalyptus* are largely conserved among species (Halliwell et al., 2025), these convergent strategies are likely underpinned by different sets of genes. The lack of shared outliers further suggests that adaptation in each pair has drawn on largely independent sources of genetic variation (whether new mutation or lineage-specific standing variation) rather than a shared pool of ancestral or introgressed alleles. This is also likely given the phylogenetic separation of our three pairs across different *Eucalyptus* sections, which limits the opportunity for shared standing variation or adaptive introgression between species. The *E. nitida*–*E. regnans* pair, which does not span a transition into semi-arid conditions, showed just as little overlap with the wet-dry pairs as the wet-dry pairs showed with one another, indicating that lineage-specificity of divergence is a general feature of genomic divergence along precipitation gradients in *Eucalyptus*, rather than in relation to an aridity threshold.

Within each species pair, outlier windows contained several genes with known roles in drought response such as *MYC2* (Abe et al., 1997), *ERD4* (Chakraborty et al., 2023), *XERICO* (Ko et al., 2006), *GA2ox2* (Z. Chen et al., 2019), and *SAG12* (Merewitz et al., 2012), which represent promising lineage-specific candidates for adaptation to aridity. The small number of genes that fell within outlier windows shared across pairs (*WAK4*, *SHT*, *CYP96A10*, and *PDCD2*) are candidates worth pursuing as potential common contributors to drought adaptation across *Eucalyptus*. More broadly, although sharing of individual loci was minimal, we found some evidence for repeatability at the level of biological function. Only one process, defence response, was significantly enriched in more than one pair, and functional profiles were otherwise largely pair-specific. This is consistent with recent work in other systems showing that the repeatability of adaptation is often more evident at the level of pathways or biological functions than at the level of individual genes for polygenic traits (James et al., 2021; Lim et al., 2019; Szukala et al., 2023; Tenaillon et al., 2012). This contrasts with hybrid and experimental-evolution systems where adaptation can be highly repeatable even at individual loci (Langdon et al., 2024; Owens et al., 2025), a predictability likely driven by the shared starting conditions and strong, large-effect selection characteristic of those systems. Our *Eucalytpus* work therefore reinforces the view that the predictability of the genetic basis of adaptation is low at the level of individual loci but can emerge at higher levels of biological organisation for complex quantitative traits (Bohutínská & Peichel, 2024; Hoitinga & Birkeland, 2025; Yeaman et al., 2018).

### The interconnection between drought and pathogen response

One of the few functional signatures shared across our species pairs was enrichment for defence response, in both the *E. dealbata*–*E. vicina* and *E. nitida*–*E. regnans* comparisons, alongside the enrichment of the *E. nitida*–*E. regnans* pair for jasmonic acid and flavonoid biosynthesis. At first glance, the emergence of pathogen-defence functions is surprising, but it aligns with previous work in plants showing that the signalling networks governing drought and pathogen responses are interconnected (Atkinson & Urwin, 2012; Fujita et al., 2006; J. Yang et al., 2019). This link is well documented in *Eucalyptus*, where drought stress increases susceptibility to fungal pathogens (Barradas et al., 2018; Hossain et al., 2019; Santos et al., 2024). Abscisic acid (ABA), the central hormone of the dehydration response, interacts extensively with the jasmonic acid (JA) and salicylic acid (SA) pathways that coordinate pathogen defence, a crosstalk also evident in the hormonal responses of *Eucalyptus* to drought and infection (Santos et al., 2024). Several of the genes we identified are involved in these pathways, most notably *MYC2,* a transcription factor induced by dehydration (Kazan & Manners, 2013). The recurrence of defence functions across our species pairs has two possible explanations. In *Eucalyptus*, aridity and biotic pressures may be correlated across the landscape, with drier environments imposing distinct pathogen or herbivore communities. Alternatively, selection on the core dehydration-response network may pleiotropically select for defence-associated genes, because the two systems share signalling machinery. Disentangling these possibilities is an important direction for future work, and will require linking patterns of genomic divergence to both biotic and abiotic variables.

### Large genomic islands of differentiation

Beyond individual outlier windows, we identified 23 broad islands of elevated differentiation, concentrated in the *E. moluccana*–*E. largiflorens* and *E. nitida*–*E. regnans* pairs. Across all three species pairs, these islands were characterised primarily by strongly reduced nucleotide diversity, accompanied by reduced or unchanged *d*_XY_ and a tendency toward reduced recombination, consistent with a substantial role of linked selection in their formation. This is similar to findings in sunflowers, where genomic islands of divergence were shown to coincide with regions of reduced recombination rather than being shaped by the geography of speciation or levels of gene flow, leading to the conclusion that genome architecture is a stronger determinant of the divergence landscape than the ecological context of divergence itself (Renaut et al., 2013). Several islands harboured drought-related genes including transcription factors central to the drought and abscisic acid (ABA) regulatory network, including *MYC2* (Kazan & Manners, 2013), VNI2 (S.-D. Yang et al., 2011), *DRIP1* (Qin et al., 2008) and *CBF4* (Haake et al., 2002). The presence of these ABA-and dehydration-responsive regulators within islands of elevated differentiation suggests that, although islands are dominated by the signature of linked selection, some may also harbour genes of adaptive relevance to aridity adaptation. This clustering of differentiation into large genomic regions mirrors findings in other systems, where adaptive variation is concentrated in large haploblocks or supergenes maintained by suppressed recombination (Küpper et al., 2016; Todesco et al., 2020).

In the three largest islands, elevated F_ST_ coincided with an extended block of high linkage disequilibrium, sharply reduced nucleotide diversity in both populations, and suppressed recombination. This combination is the classic signature of a region of restricted recombination, such as a chromosomal inversion or a centromeric region (K. Huang & Rieseberg, 2020). Chromosomal inversions are well established as drivers of local adaptation across diverse taxa including *Mimulus* (Lowry & Willis, 2010), *Heliconius* (Joron et al., 2011), *Anopheles* (Ayala et al., 2014), and deer mice (Harringmeyer & Hoekstra, 2022). By suppressing recombination in heterozygotes, they trap locally adapted alleles together in the face of gene flow and allow differentiation to accumulate (Kirkpatrick & Barton, 2006). In *Eucalyptus*, inversions have been linked to both climate adaptation and divergence between species (Ferguson et al., 2024; Zhuang et al., 2026), although comparative genomic work suggests that fixed inversions between *Eucalyptus* genomes are relatively rare and contribute less to divergence than other classes of rearrangements (Ferguson et al., 2024). Future work further characterising the genomic landscape of structural variation will help elucidate the mechanisms driving recombination suppression in these large regions of elevated differentiation.

### Interpreting genomic landscapes of diversity and divergence

Our work identified genomic regions associated with the transition into drier environments; however, several caveats should be considered when interpreting our results. First, across Australia, many *Eucalyptus* species exhibit pervasive historical and contemporary gene flow (Fahey et al., 2022; Flores-Rentería et al., 2017; Jones et al., 2016), which influences how signatures of natural selection persist in the genome. Introgression can homogenise large portions of the genome and obscure shared adaptive signals, reducing the repeatability of individual loci across lineages. Furthermore, given that climate adaptation in *Eucalyptus* is polygenic, our outlier approach likely captures only a subset of loci contributing to adaptive divergence and preferentially identifies regions with larger allele frequency differentiation. Many adaptive alleles may instead experience subtle frequency shifts and therefore remain undetected by outlier scans (Yeaman, 2015). Despite this limitation, the genes identified here represent strong candidates for the repeated adaptation to drier environments. Finally, although we have explored replicated evolution in relation to transitions to drier environments, convergence can also occur due to unmeasured factors that are correlated with the environment (Rellstab et al., 2015). Future work directly correlating genomic variation to variation in climatic variables is key to disentangling this.

Overall, our work shows that the transition into drier environments in *Eucalyptus* is shaped by pervasive linked selection that produces broadly similar diversity and divergence landscapes across lineages, yet the adaptive responses are largely non-convergent across lineages. Our results suggest that the genetic basis of complex polygenic traits, such as aridity adaptation, is largely unpredictable at the level of individual genes yet more repeatable at the level of biological function. This has implications for how we understand the predictability of adaptation under a drying climate, as the precise genes underpinning aridity adaptation in one *Eucalyptus* species cannot be assumed to predict those in another.

## ACKNOWLEDGEMENTS

We acknowledge the Turrbal, Yuggera, Darug, Dharawal, Gadigal, Gundungurra, Wiradjuri, Palawa, Gadigal and Wangal peoples, the traditional owners of the land on which this work was undertaken. We acknowledge the Darug, Wiradjuri, Kamilaroi, Lairmairrener, Nuenonne and Tharawal peoples, the traditional owners of the land on which the samples were collected. We pay our respects to Elders past, present and emerging. We are grateful to Stuart Allen and Rachael Gallagher for providing species climate data, and we thank Dean Nicolle for confirming species identifications. We acknowledge the use of Claude Opus v4.8 to assist with grammatical edits to the main text and code development for data visualisation. This research was supported by a Kickstart Grant to CI Maddie James through the Australian Research Council Centre of Excellence for Plant Success in Nature and Agriculture (CE200100015).

## AUTHOR CONTRIBUTIONS

**MEJ:** Conceptualisation (lead), Funding Acquisition (lead), Supervision (lead), Investigation (lead), Data Curation (lead), Formal Analysis (lead), Visualisation (lead), Writing – Original Draft (lead), Writing – Review & Editing (lead), Project Admin (lead). **TJB:** Conceptualisation (supporting), Funding Acquisition (supporting), Investigation (supporting), Visualisation (supporting), Writing – Review & Editing (supporting). **JDM:** Conceptualisation (supporting), Funding Acquisition (supporting), Formal Analysis (supporting), Writing – Review & Editing (supporting). **BH:** Conceptualisation (supporting), Funding Acquisition (supporting), Writing – Review & Editing (supporting). **BH, IJW and DOB:** Supervision (supporting), Writing – Review & Editing (supporting).

## SUPPLEMENTARY FIGURES

**Figure S1.**
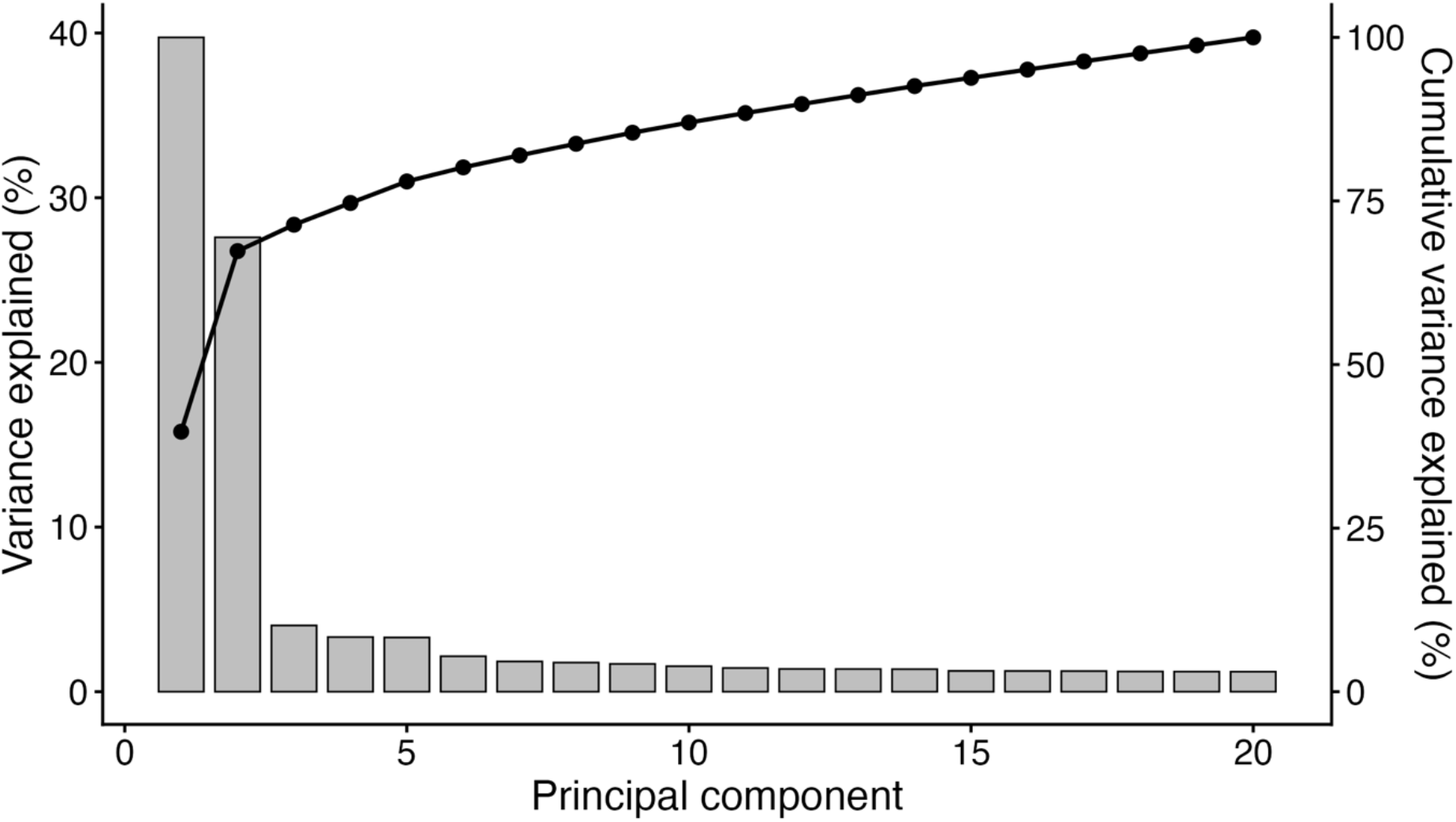
Principle Components Analysis (PCA) scree plot. Variance explained and cumulative variance for the first 20 PCs using 428,330 unlinked SNPs for the nine *Eucalyptus* species.

**Figure S2.**
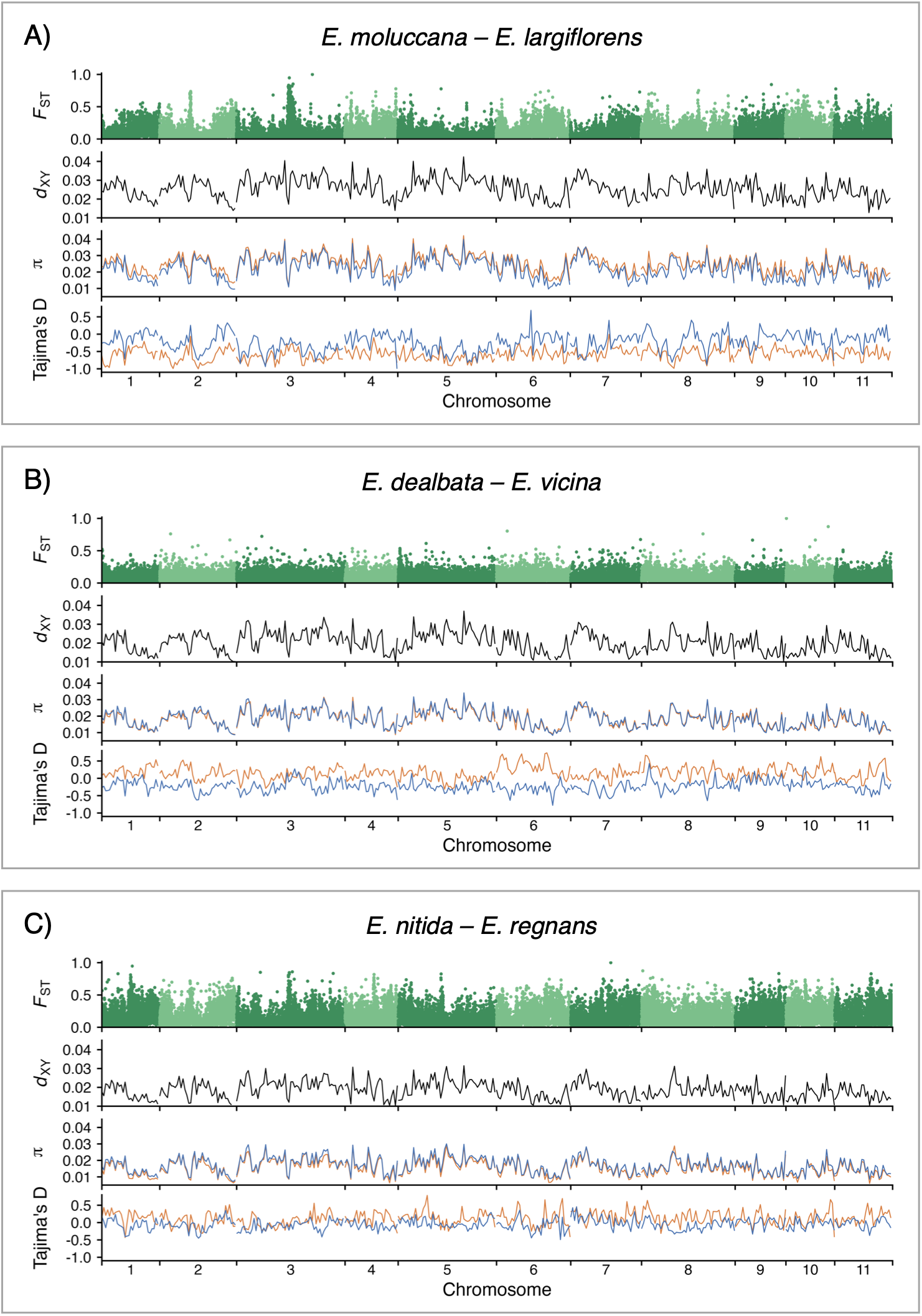
Genome-wide landscapes of diversity and divergence. Population genetic statistics across the 11 chromosomes for A) *Eucalyptus moluccana* (wet) vs *E. largiflorens* (dry), B) *E. dealbata* (wet) vs *E. vicina* (dry), and C) *E. nitida* (wetter) vs *E. regnans* (wet) calculated in 20kB non-overlapping sliding windows. Top panel: genome-wide distribution of F_ST_ values showing levels of genetic differentiation between species. Second panel: *d*_XY_ values along the genome, representing absolute divergence between species. Third panel: nucleotide diversity (π) for wetter (blue) and drier (orange) species along the genome, showing intraspecific genetic variation. Bottom panel: Tajima’s D statistics, where positive values indicate an excess of intermediate-frequency alleles (consistent with balancing selection or population contraction), and negative values indicate an excess of rare alleles (consistent with selective sweeps, purifying selection or population expansion). *d*_XY_, π and Tajima’s D plots were smoothed by calculating mean values across 1.5Mb bins and connecting adjacent bins with lines.

**Figure S3.**
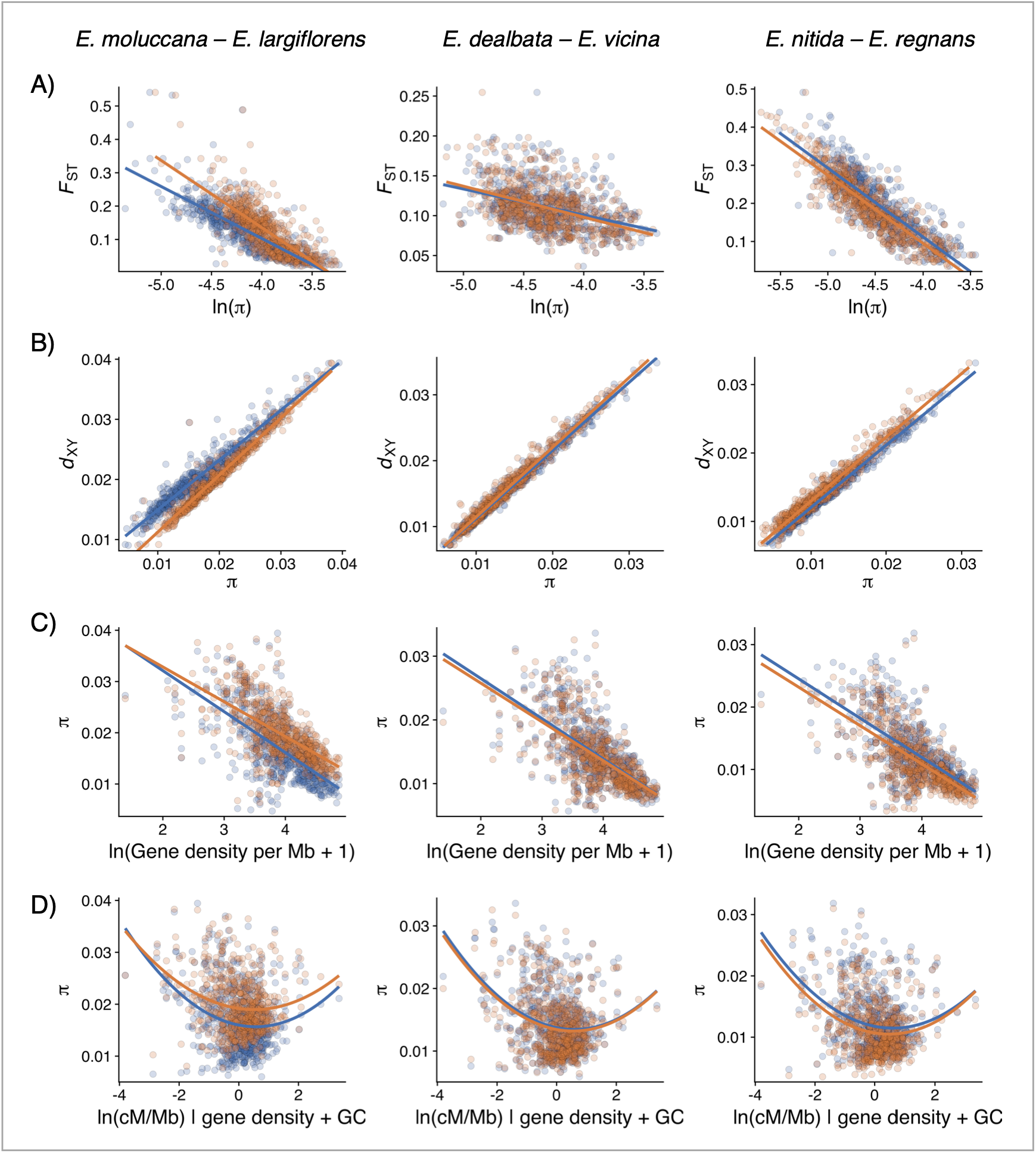
Evidence for linked selection. Scatterplots of pairwise relationships across the genome for *E. moluccana*–*E. largiflorens*, *E. dealbata*–*E. vicina* and *E. nitida*–*E. regnans* for the wet (blue) and dry (orange) species for A) F_ST_ vs π, B) *d*_XY_ vs π, C) Mean π vs gene density per Mb, and D) π vs recombination rate, after accounting for gene density and GC content. Each point represents a 1Mb non-overlapping genomic window; trend lines show linear regressions for visualisation.

**Figure S4.**
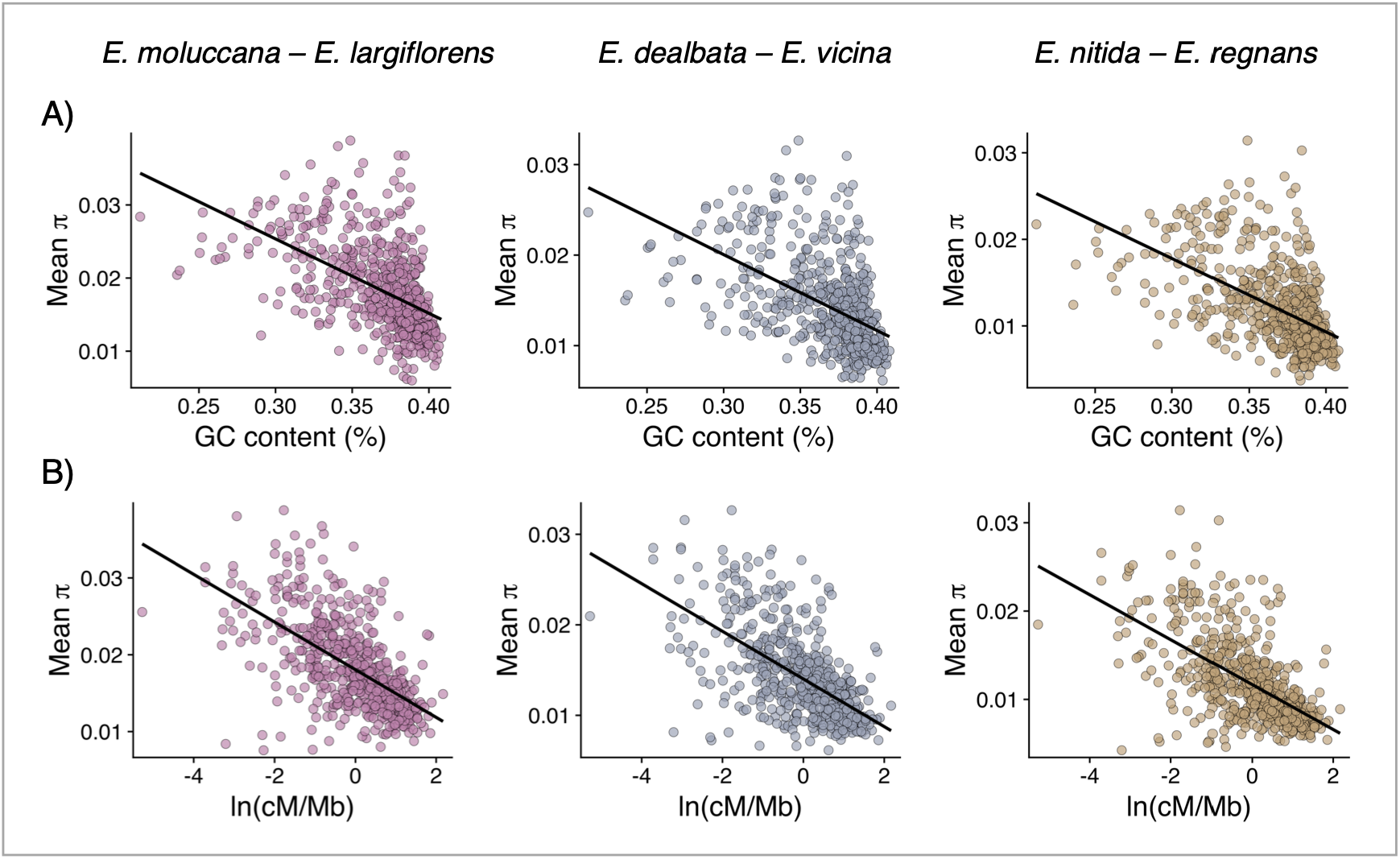
Correlations between π and genomic features. Scatterplots of pairwise relationships across the genome for *E. moluccana*–*E. largiflorens* (pink), *E. dealbata*–*E. vicina* (slate) and *E. nitida*–*E. regnans* (brown) for A) Mean π vs GC content, and B) Mean π vs recombination rate. Each point represents a 1Mb non-overlapping genomic window; trend lines show linear regressions for visualisation. See Table S2 for Pearson’s and Spearman’s correlation coefficients and associated P-values.

**Figure S5.**
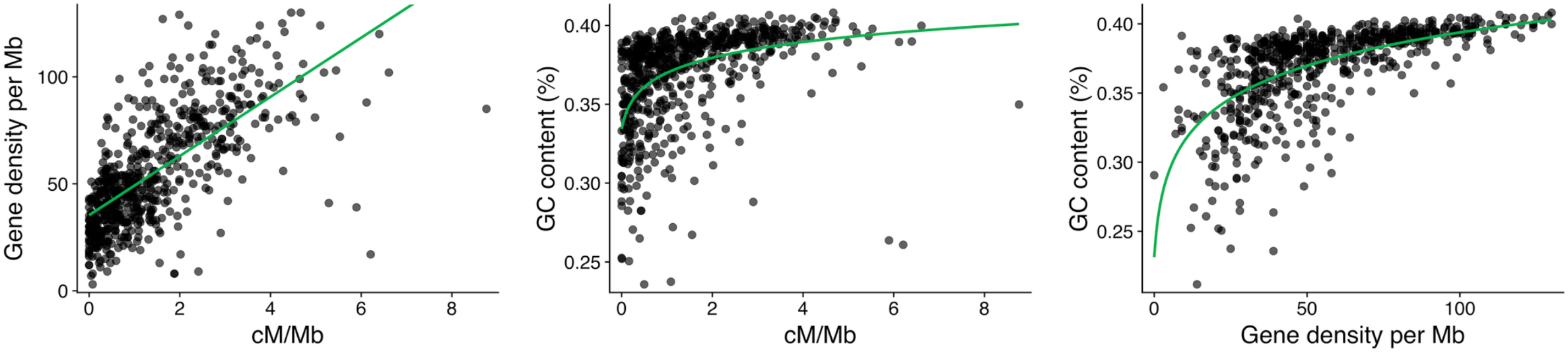
Relationships among genome features. Scatterplots of pairwise relationships for gene density p (cM/Mb) (left plot), GC content vs recombination rate (cM/Mb) (middle plot), and GC content vs gene densi represents a 1Mb non-overlapping genomic window; trend lines show fitted curves for visualisation (left: linear; middle and right: logarithmic).

**Figure S6.**
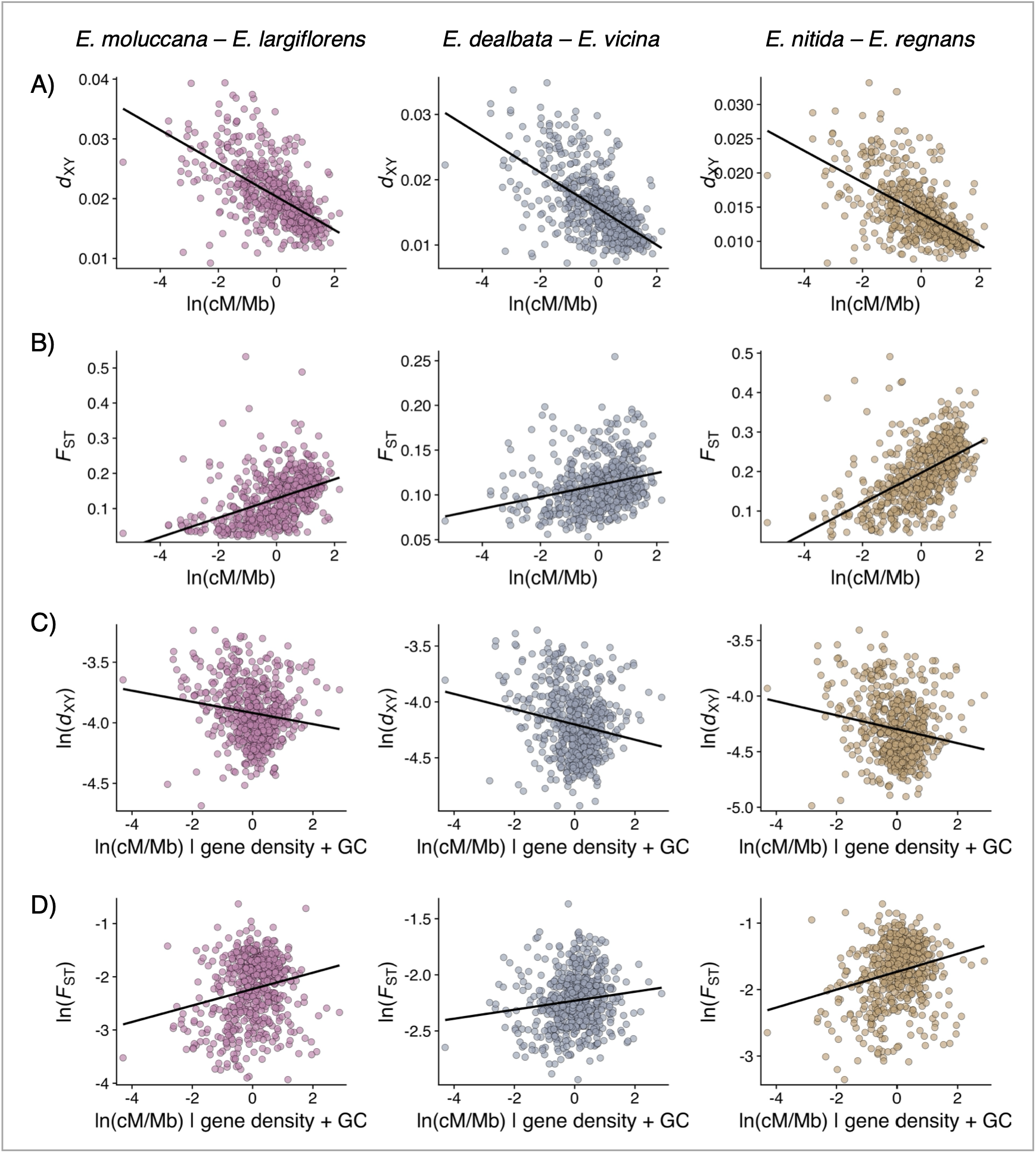
Evidence for linked selection. Scatterplots of pairwise relationships across the genome for *E. moluccana–E. largiflorens* (pink), *E. dealbata–E. vicina* (slate), and *E. nitida–E. regnans* (brown). A) *d*_XY_ vs recombination rate and B) F_ST_ vs recombination rate. C) *d*_XY_ and D) F_ST v_s recombination rate, after accounting for gene density and GC content. Each point represents a 1Mb non-overlapping genomic window; trend lines show linear regressions for visualisation. See Table S2 for Pearson’s and Spearman’s correlation coefficients and associated P-values.

**Figure S7.**
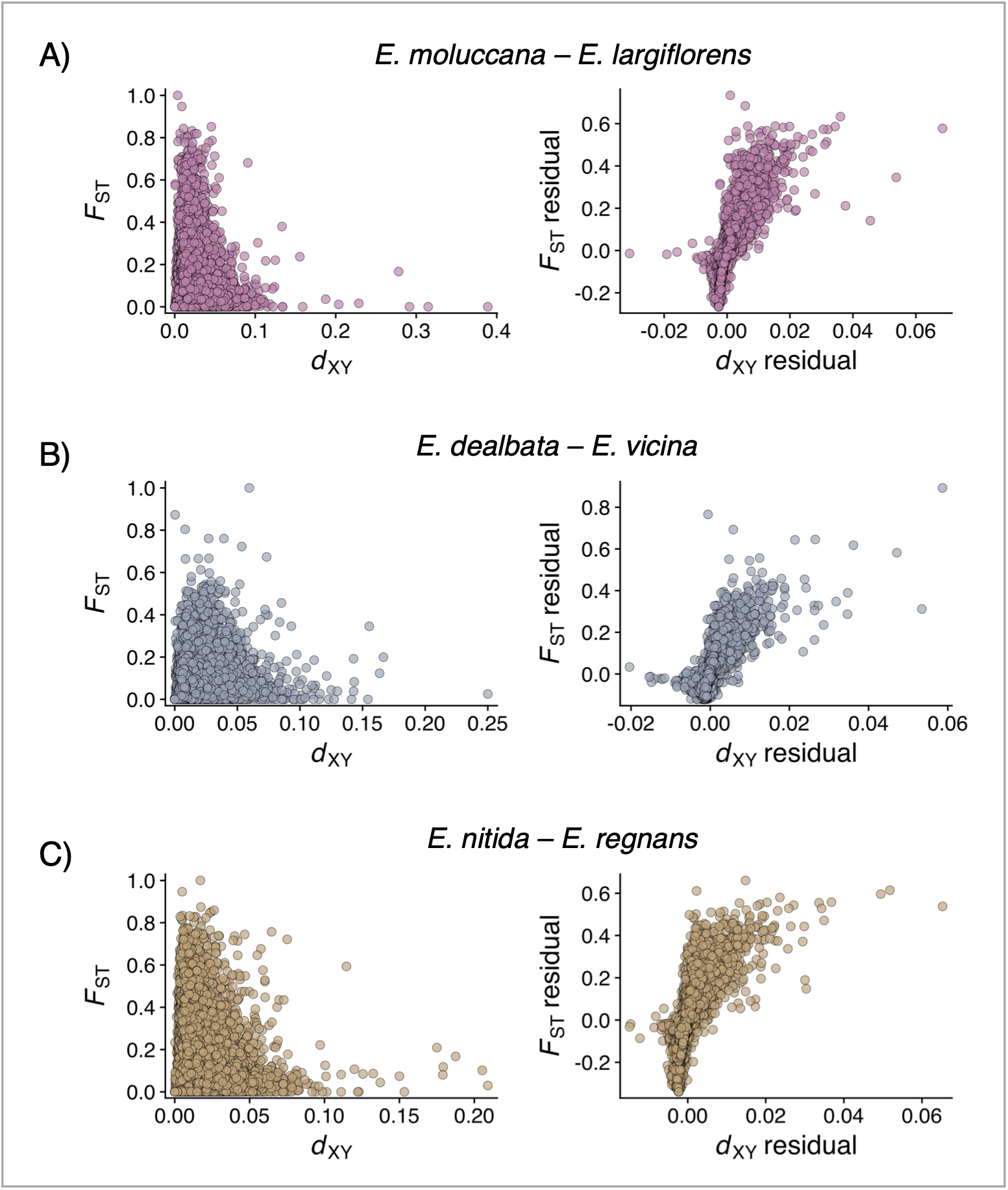
Correlations between F_ST_ and *d*_XY_. Scatterplots of pairwise relationships for F_ST_ vs *d*_XY_ and F_ST_ residuals vs *d*_XY_ residuals (after taking into account π for A) *E. moluccana–E. largiflorens* (pink), B) *E. dealbata–E. vicina* (slate), and C) *E. nitida–E. regnans* (brown). Once background diversity levels are controlled for, regions with elevated absolute divergence also show elevated relative differentiation.

**Figure S8.**
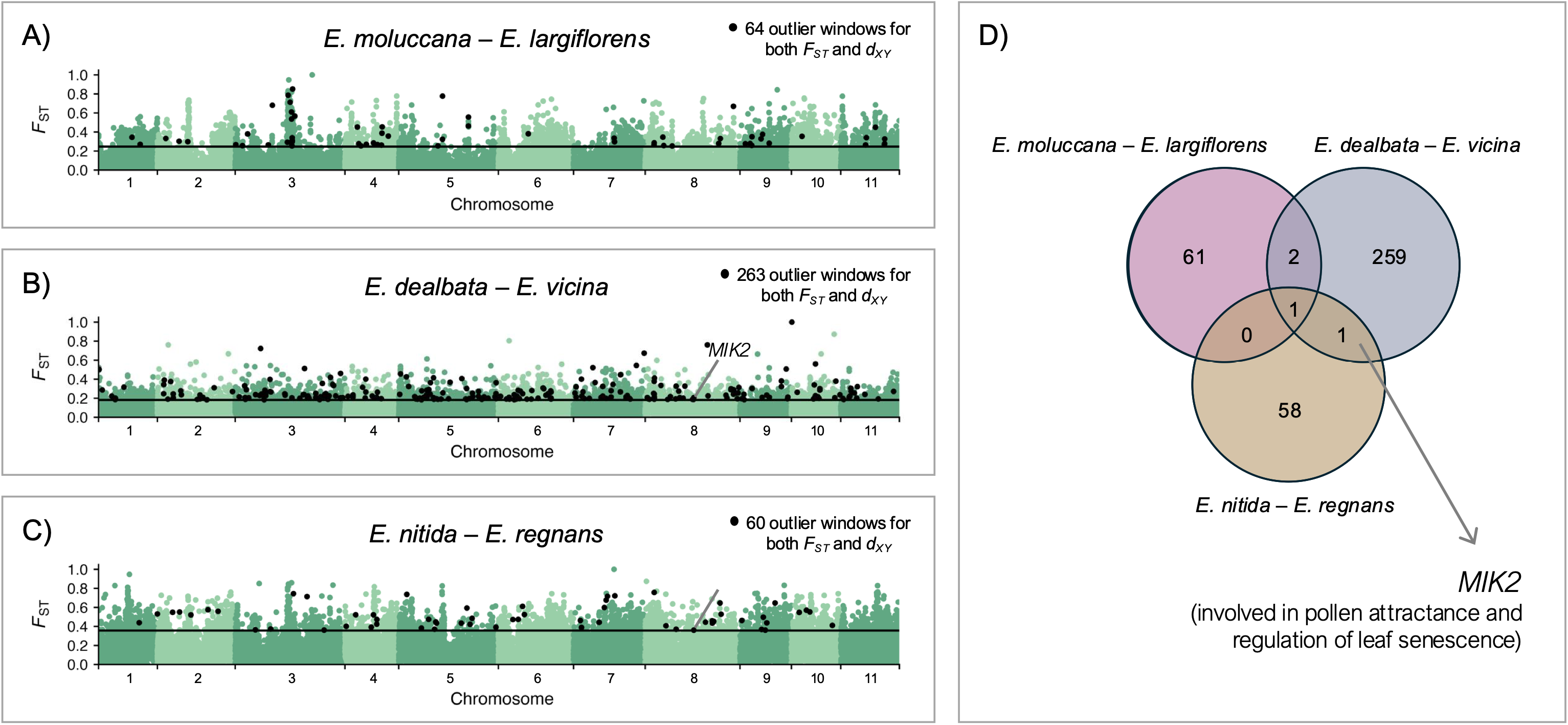
Outlier analysis. F_ST_ across the 11 *Eucalyptus* chromosomes for A) *E. moluccana*–*E. largifloren E. nitida*–*E. regnans*. The black horizonal lines indicate the significance threshold (top 10% of genomic wind non-overlapping window. Points coloured black represent outlier windows found in the top 10% of both F_ST_ windows across the three species pairs. Arrows and gene descriptions represent genes that reside in any share

**Figure S9.**
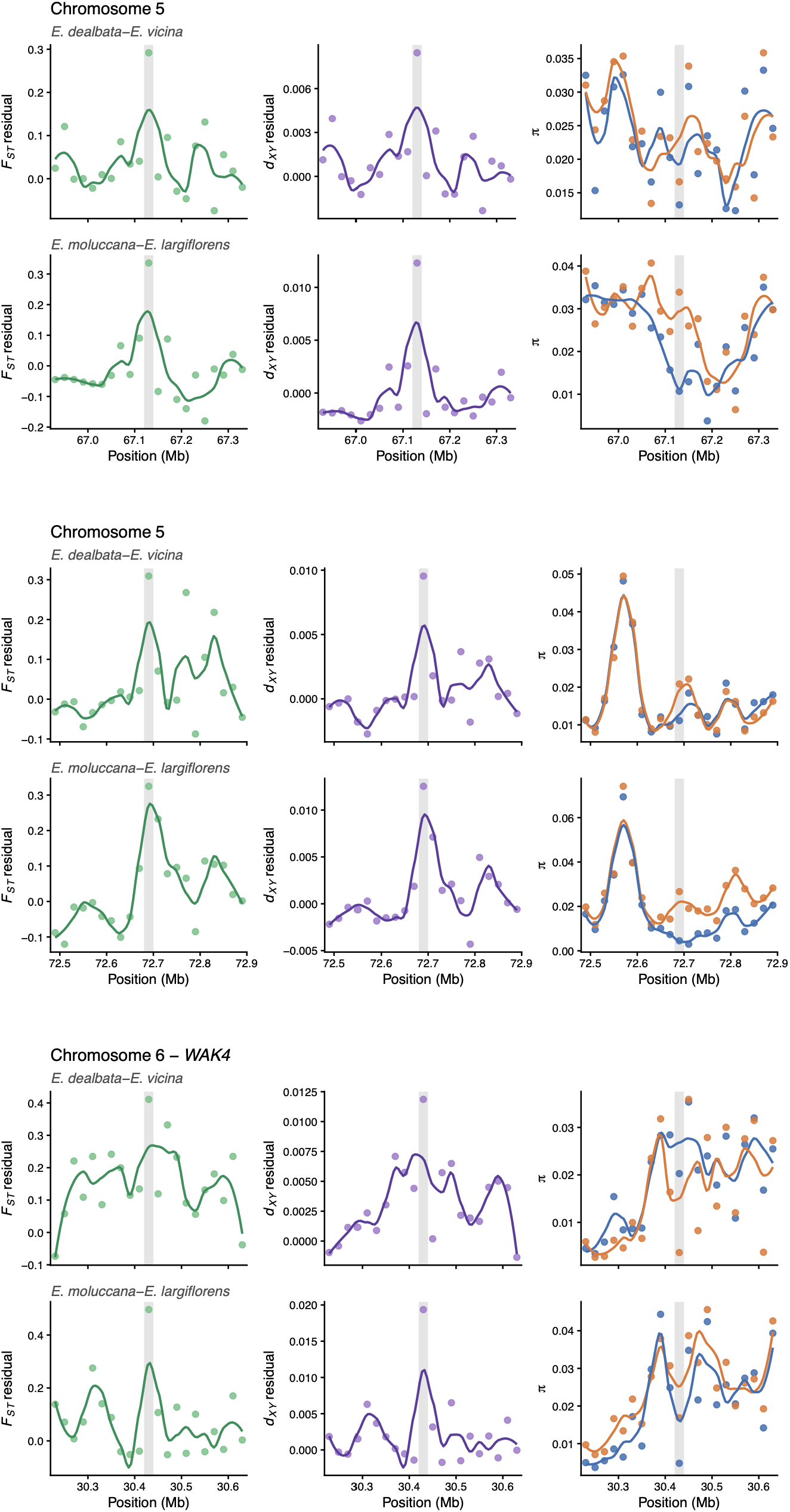
Shared outlier windows between *E. moluccana*–*E. largiflorens* and *E. dealbata*–*E. vicina*. Zoom in of surrounding regions of shared outlier windows. Each dot represents a 20kB non-overlapping window. Grey shading represents the outlier window.

**Figure S10.**
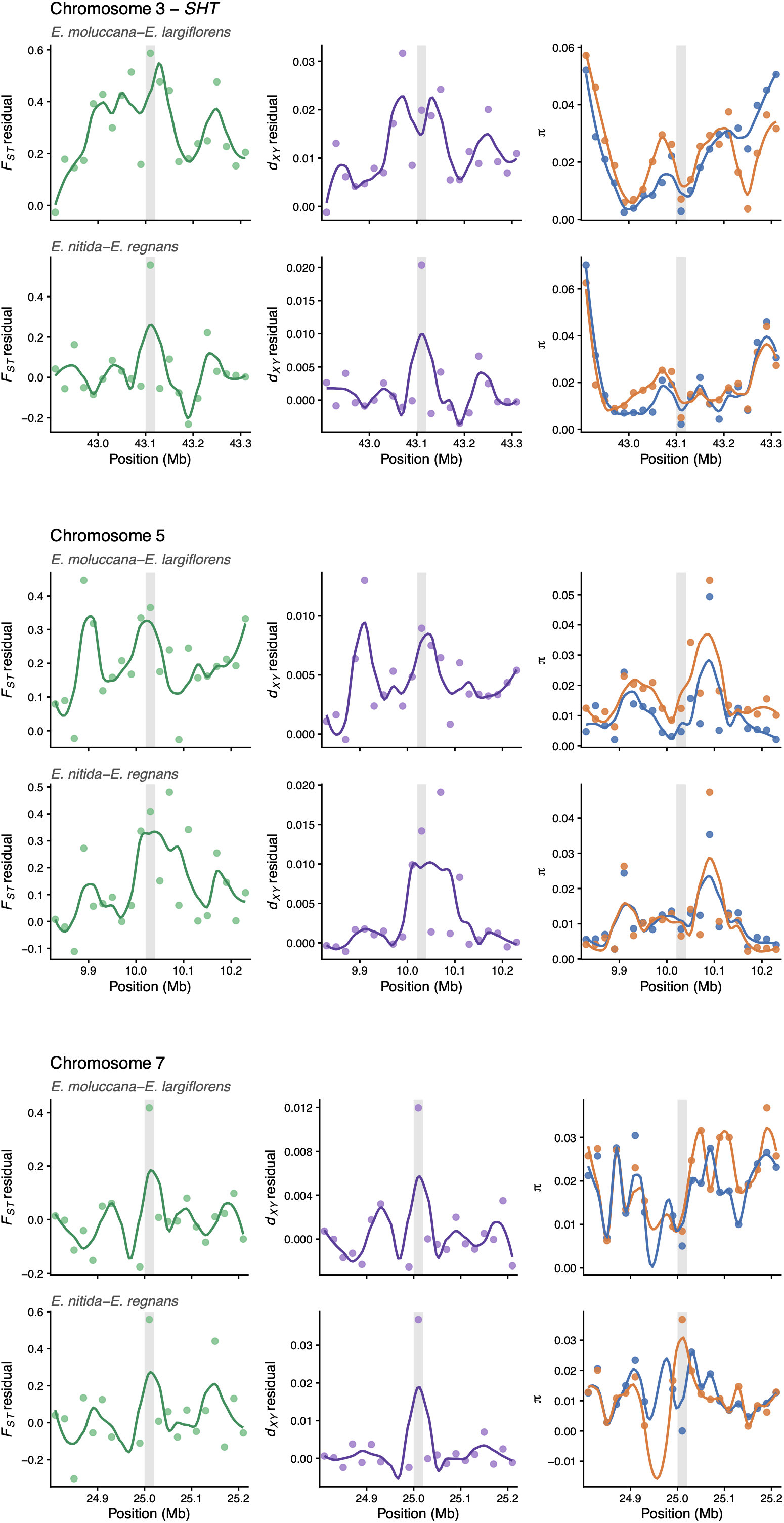
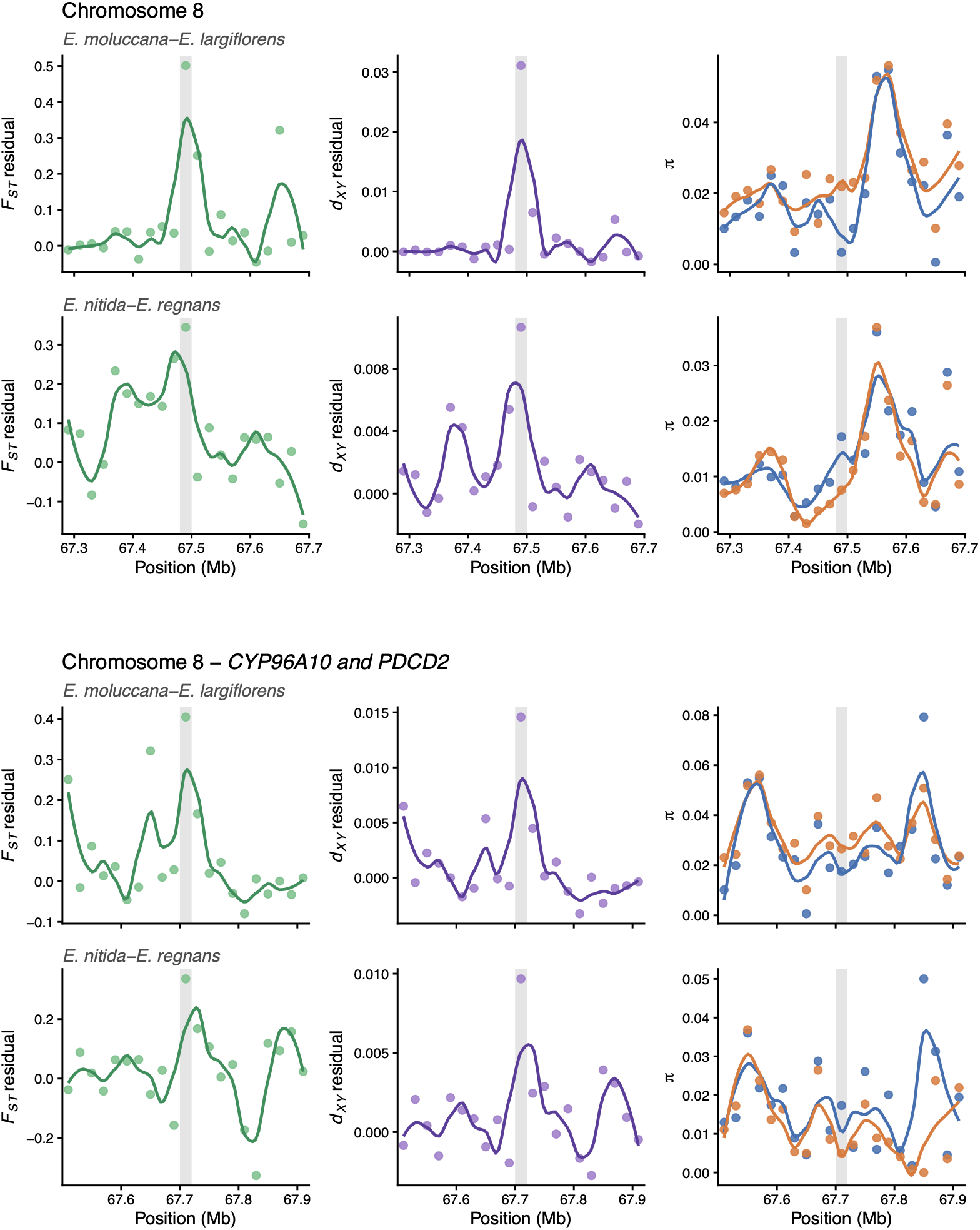
Shared outlier windows between *E. moluccana*–*E. largiflorens* and *E. nitida*–*E. regnans*. Zoom in of surrounding regions of shared outlier windows. Each dot represents a 20kB non-overlapping window. Grey shading represents the outlier window.

**Figure S11.**
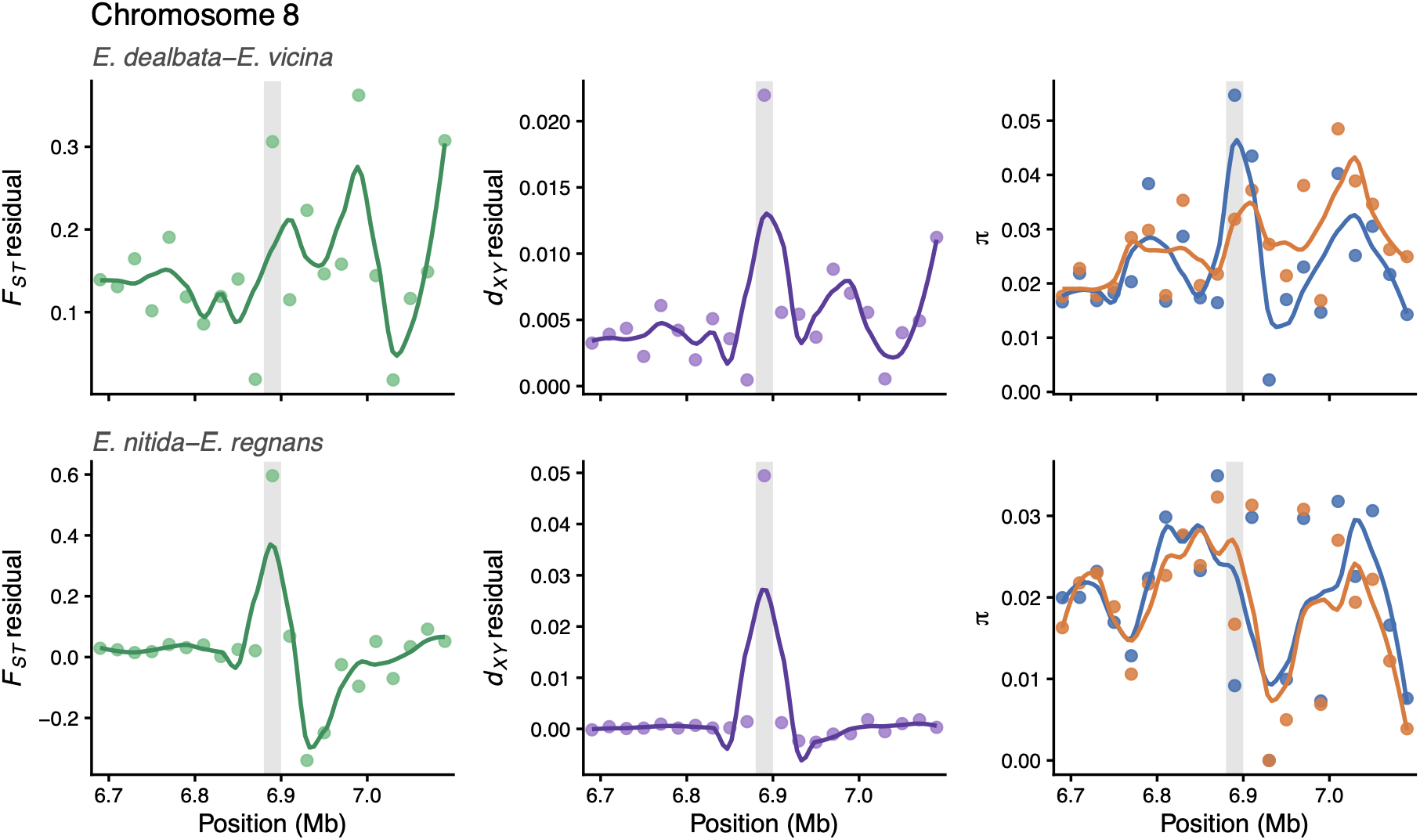
Shared outlier window between *E. dealbata*–*E. vicina* and *E. nitida*–*E. regnans*. Zoom in of surrounding regions of shared outlier windows. Each dot represents a 20kB non-overlapping window. Grey shading represents the outlier window.

**Figure S12.**
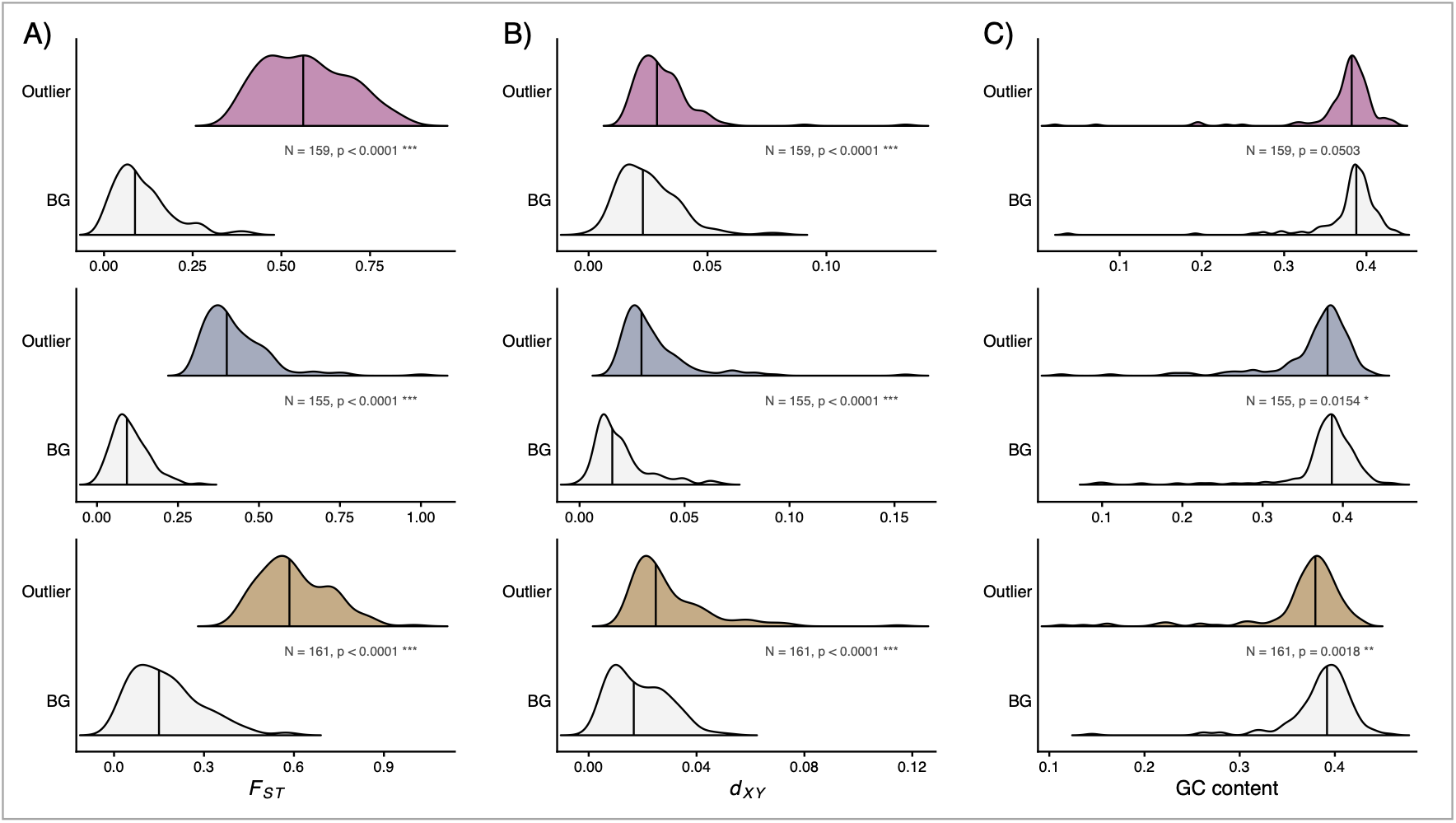
Associations of outliers with genome features. Half-violin plots for outlier windows (top, coloured) and a random sample of background windows (BG, bottom, grey) for A) F_ST_, B) *d*_XY_ residuals, and C) GC content. Vertical bars represent the median of each distribution. Text is a permutation test comparing outlier and background windows, in which the observed median of the outlier windows was compared against a null distribution generated by drawing 10,000 random samples of background windows matched to the number of outlier windows (N = number of outlier windows and two-sided empirical P-value; * p < 0.05, ** p < 0.01, *** p < 0.001). Top row: *E. moluccana*–*E. largiflorens* (pink); middle row: *E. dealbata*–*E. vicina* (slate); bottom row: *E. nitida*–*E. regnans* (brown). Outlier windows were defined as the top 1% of windows for both F_ST_ and *d*_XY_ residuals.

**Figure S13.**
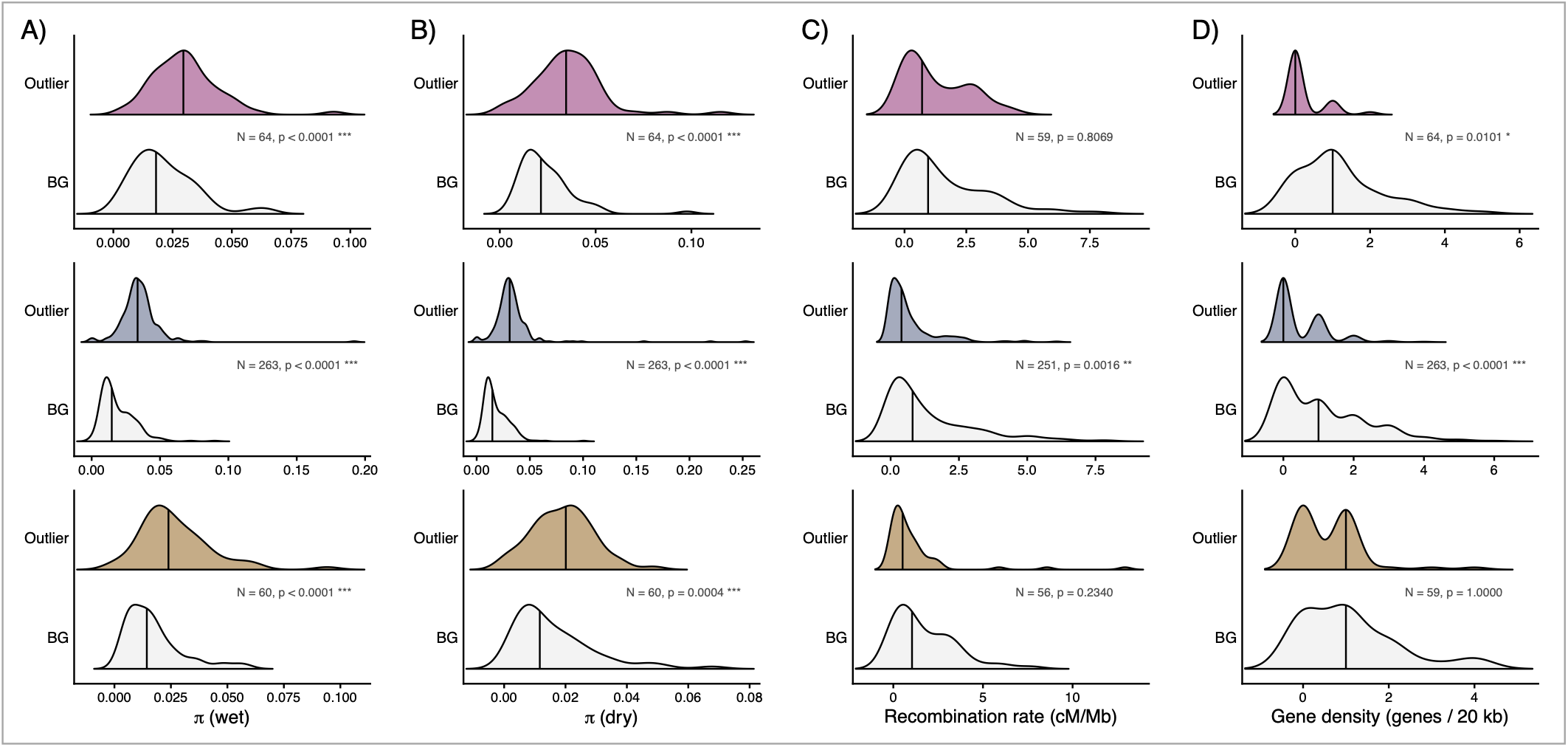
Associations of outliers with genome features – top 10% F_ST_ and *d*_XY_ outlier approach. Half-violin plots for outlier windows (top, coloured) and a random sample of background windows (BG, bottom, grey) for A) nucleotide diversity (π) in the wet-adapted species, B) π in the dry-adapted species, C) recombination rate (cM/Mb), and D) gene density (genes per 20 kb). Vertical bars represent the median of each distribution. Text is a permutation test comparing outlier and background windows, in which the observed median of the outlier windows was compared against a null distribution generated by drawing 10,000 random samples of background windows matched to the number of outlier windows (N = number of outlier windows and two-sided empirical P-value; * p < 0.05, ** p < 0.01, *** p < 0.001). Top row: *E. moluccana*–*E. largiflorens* (pink); middle row: *E. dealbata*–*E. vicina* (slate); bottom row: *E. nitida*–*E. regnans* (brown). Outlier windows were defined as the top 10% of windows for both F_ST_ and *d*_XY_.

**Figure S14.**
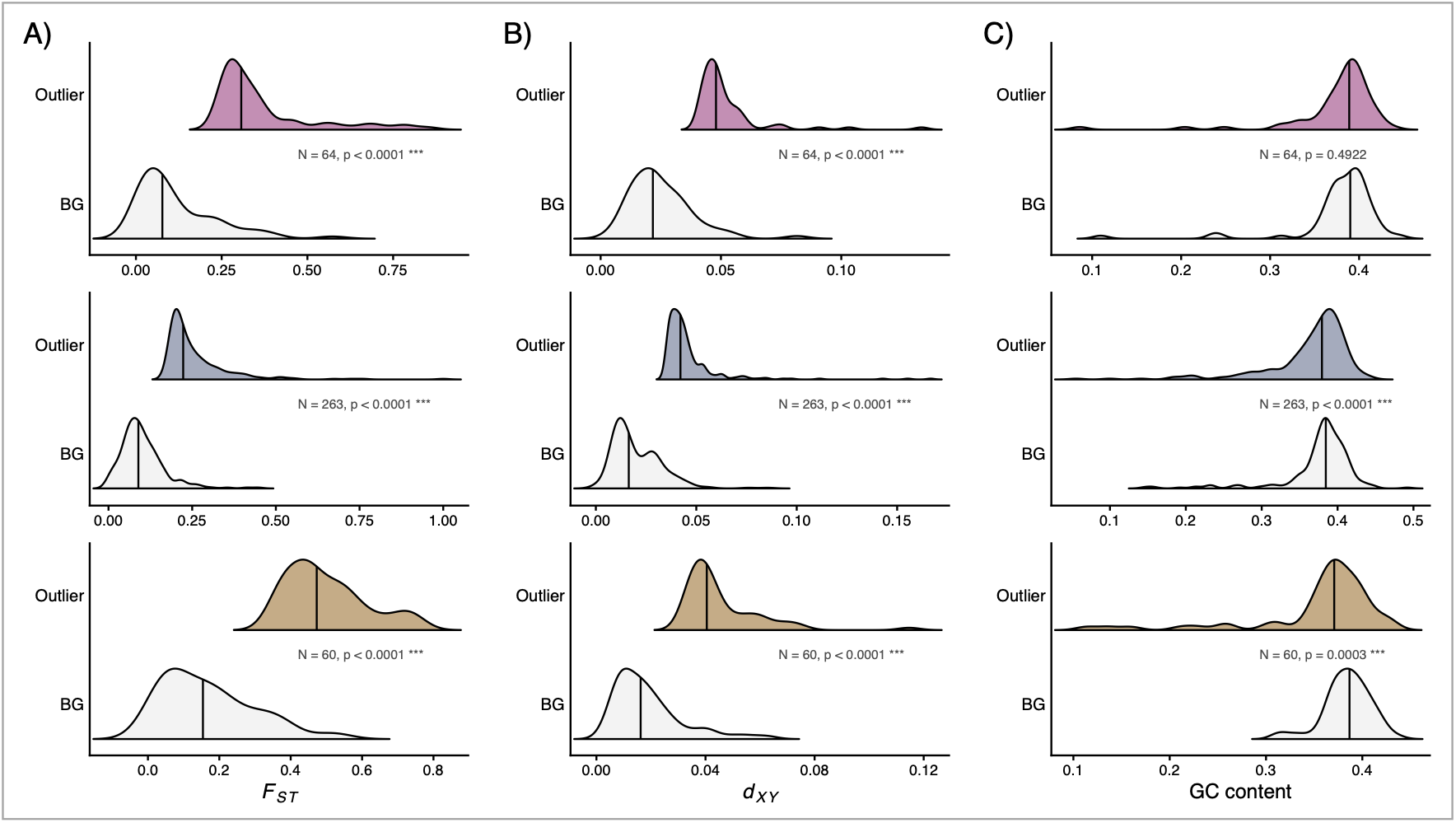
Associations of outliers with genome features – top 10% FST and *d*XY outlier approach. Half-violin plots for outlier windows (top, coloured) and a random sample of background windows (BG, bottom, grey) for A) FST, B) *d*XY residuals, and C) GC content. Vertical bars represent the median of each distribution. Text is a permutation test comparing outlier and background windows, in which the observed median of the outlier windows was compared against a null distribution generated by drawing 10,000 random samples of background windows matched to the number of outlier windows (N = number of outlier windows and two-sided empirical P-value; * p < 0.05, ** p < 0.01, *** p < 0.001). Top row: *E. moluccana*–*E. largiflorens* (pink); middldee row: *E. dealbata*–*E. vicina* (slate); bottom row: *E. nitida*–*E. regnans* (brown). Outlier windows were defined as the top 10% of windows for both FST and *d*XY.

## SUPPLEMENTARY TABLE CAPTIONS

**Table S1**. **Sampling locations and climate information of the nine *Eucalyptus* species.** Mean annual precipitation (MAP, mm), mean annual temperature (MAT, °C) and moisture index (MI, unitless) describe the mean climate conditions across each species current-day geographic distribution. Pair ID identifies matched species pairs used in comparative analyses (NA indicates an outgroup species), and Pair Type denotes whether species belong to the main paired design or additional/outgroup sampling. Climate Category indicates whether each species represents the wetter or drier member of its pair (or “wetter” in cases where both species occupy relatively wet environments).

**Table S2. Correlations of genomic diversity and divergence statistics.** Pearson’s *r* and Spearman’s *ρ* correlations, and linear model statistics calculated across 20kb or 1Mb non-overlapping windows for six *Eucalytpus* species.

**Table S3. *E. moluccana* – *E. largiflorens* outlier windows containing genes, detected using the residual-based outlier approach.** Outlier windows were defined as those in the top 1% for both F_ST_ residuals and *d*_XY_ residuals.

**Table S4. *E. dealbata* – *E. vicina* outlier windows containing genes, detected using the residual-based outlier approach.** Outlier windows were defined as those in the top 1% for both F_ST_ residuals and *d*_XY_ residuals.

**Table S5. *E. nitida* – *E. regnans* outlier windows containing genes, detected using the residual-based outlier approach.** Outlier windows were defined as those in the top 1% for both F_ST_ residuals and *d*_XY_ residuals.

**Table S6. Gene ontology enrichment analysis for the residual-based outlier approach.** Outlier windows were defined as those in the top 1% for both F_ST_ and *d*_XY_ residuals.

**Table S7. *E. moluccana* – *E. largiflorens* outlier windows containing genes, detected using the top 10% outlier approach.** Outlier windows were defined as those in the top 10% for both F_ST_ and *d*_XY_ raw values.

**Table S8. *E. dealbata* – *E. vicina* outlier windows containing genes, detected using the top 10% outlier approach.** Outlier windows were defined as those in the top 10% for both F_ST_ and *d*_XY_ raw values.

**Table S9. *E. nitida* – *E. regnans* outlier windows containing genes, detected using the top 10% outlier approach.** Outlier windows were defined as those in the top 10% for both F_ST_ and *d*_XY_ raw values.

**Table S10. Gene ontology enrichment analysis for the top 10% outlier approach.** Outlier windows were defined as those in the top 10% for both F_ST_ and *d*_XY_ raw values.

**Table S11. Genomic islands of differentiation.** Islands of differentiation and summary statistics of each island across the three species pairs.

**Table S12. Gene ontology enrichment analysis for the genomic islands of differentiation.** Enrichment analysis was undertaken for each pair with all genes located within the islands of differentiation.

**Table S13. Islands of differentiation genes.** Summary of the genes, their functions and locations within the islands of differentiation for each pair.

